# Distinct A*β*40 and A*β*42 co-assembly pathways with free lipids: Experiment-driven MD study^†^

**DOI:** 10.64898/2026.09.28.755027

**Authors:** Rachit Pandey, Tarek Olabi, Myungwoon Lee, Brigita Urbanc

## Abstract

Amyloid *β*-protein (A*β*), which is known for its role in Alzheimer’s disease (AD), is also involved in several physiological functions. Of the two predominant isoforms, A*β*40 and A*β*42, the latter is more aggregation-prone and more active in health and disease. In the brain, A*β* interacts with free lipids, which may affect its assembly pathways. Here, thioflavin T (ThT) fluorescence spectroscopy is combined with molecular dynamics (MD) to investigate A*β*40 and A*β*42 co-assembly with free dimyristoylphosphatidylcholine (DMPC) and DMPC/cholesterol (CHOL) lipids. ThT measurements at several protein-to-lipid (PL) ratios show isoform- and lipid composition-dependent kinetics of A*β* fibril formation with clear isoform-dependent lipid effects at a PL ratio of 1:5. Early co-assembly stages of six A*β*40/A*β*42 with DMPC and DMPC/CHOL lipids at this PL ratio are then examined by MD. MD results indicate that lipids alter the secondary structure more and interact more avidly with A*β*40 than with A*β*42. In turn, A*β*40 increases the planar order of lipid bicelles more than A*β*42. Unlike A*β*40, A*β*42 decreases the nematic order of CHOL. In DMPC/CHOL lipids, A*β*42 but not A*β*40 exhibits two A*β*-lipid interaction modes associated with mutually exclusive A*β*42-lipid and A*β*42-A*β*42 interactions. Our MD findings provide a plausible explanation for an experimentally observed increase in the lag time for A*β*40 fibril formation in DMPC lipids and even more so in DMPC/CHOL lipids that contrasts with much smaller effects of lipids on A*β*42 fibril formation and offer additional atomistic-level insights into A*β* co-assembly with lipids amenable to experimental testing.

## 1 Introduction

Amyloid *β*-protein (A*β*) is notorious for its role in Alzheimer’s disease (AD), which is a progressive neurodegenerative disorder and the leading cause of dementia worldwide ^1^. The amyloid cascade hypothesis, which initially posited that amyloid-*β* (A*β*) aggregation into fibrils and subsequent accumulation into amyloid plaques is the basis of AD pathogenesis, has been later revised to acknowledge a critical role of soluble A*β* oligomers as the proximate neurotoxic species in AD ^2,3^. Of all naturally-occurring A*β* isoforms, A*β*40 and A*β*42 are the two predominant forms and differ only by the two additional residues, I41 and A42, at the C-terminal of A*β*42. A*β*42 is generally considered more neurotoxic ^4^, more aggregation-prone, and exhibits a greater propensity to form larger oligomeric assemblies than A*β*40^5^. A*β*42 is also one of the most effective biomarkers for AD ^6^. The lead hypothesis of how A*β* triggers AD pathology posits that A*β* oligomers interact with cellular membranes and disrupt calcium homeostasis, which leads to cell dysfunction ^7–14^. Several mechanisms have been proposed for A*β* oligomer-mediated membrane disruption, including carpeting of the membrane surface, insertion into the lipid bilayer, and extraction of lipids through a detergent-like mechanism ^15^.

Full-length, 37-43 amino acid residues-long A*β* peptides are produced throughout lifetime from the amyloid precursor protein (APP) through sequential *β*- and *γ*-secretase cleavages. Substantial evidence shows that A*β* is not just a waste byproduct of APP processing, but may be involved in several important physiological functions by participating in the antimicrobial defense in the brain, sealing leaks in the blood-brain barrier, recovering from brain injury, and regulating activity of hippocampal synapses ^16^. Notably, White and collaborators reported that the two C-terminal residues in A*β*42 are required for the antiviral and antibacterial activity, indicating that physiological activity of A*β* is also isoform dependent ^17^. Moreover, antimicrobial function of A*β* involves formation of oligomers and fibrils that interact with viral or bacterial membranes and effectively disrupt their microbial activity ^18^. Thus, A*β* exerts its physiological or toxic effects through interactions with lipids.

A*β* is an intrinsically disordered protein (IDP), which makes it challenging to study experimentally or computationally even in the absence of lipids. To efficiently sample the free energy landscape of A*β* monomer and oligomer conformations in computer simulations, enhanced sampling techniques or multiple replica MD trajectories need to be used due to large trajectory-to-trajectory variability associated with IDP conformational dynamics ^19–21^. From a biophysical point of view, different folding and assembly pathways reported for the two predominant isoforms, A*β*40 and A*β*42, in experimental and computational studies ^5,22–27^, are intriguing considering the 95% sequence similarity. In line with these findings, a recent computational comparison of the physiologically relevant C-terminal isoforms A*β* 38, A*β*40, A*β*42, and A*β*43 revealed that the C-terminal length strongly influences oligomer size distribution and conformational ensembles of the resulting oligomers ^28^.

In addition to intrinsic sequence-dependent effects, interactions with lipids can substantially alter A*β* conformational dynamics and assembly, while A*β*, in turn, can perturb the organization and properties of surrounding lipids. Molecular dynamics (MD) simulations have revealed many aspects of A*β*-lipid interactions, however, most MD studies on A*β*-lipid interactions have examined A*β* interacting with a preformed lipid bilayer, whereby A*β* monomers or oligomers are initially placed into a vicinity or inserted into a lipid bilayer ^29–35^. In this manner, the effect of lipid composition on A*β*-lipid interactions has been examined by Brown and Bevan, who reported that A*β*42 tetramers interact differently with pure POPC and cholesterol (CHOL)-rich lipid bilayers, resulting in different degrees of membrane perturbation ^31^. Lipid composition has been shown to also influence the structure of A*β* itself, as Fatafta and collaborators found that the coil-to-*β*-sheet transition observed for A*β*42 dimers in aqueous solution is suppressed in the presence of a neuronal membrane-mimicking lipid bilayer through preferential interactions of dimers with gangliosides (GM1) ^33^. Similarly, varying the CHOL concentration in a pre-formed lipid bilayer has been shown to alter A*β*42 dimer stability and peptide-lipid interactions ^34^. Together, these studies demonstrate that A*β* structure, assembly state, and lipid composition clearly affect A*β* interactions with lipid bilayers.

Significantly fewer MD studies have thus far examined A*β* co-assembly with free lipids, where both biomolecules actively participate in the assembly process. While not examining exactly A*β*-lipid co-assembly, Fatafta and collaborators showed that association of A*β*42 monomers with a small number of free phospholipids alters peptide conformation and subsequently affects its interaction with a preformed lipid bilayer ^36^. The importance of a specific A*β* isoform in selection of A*β*-lipid modes of interaction has been elucidated by Chakravorty and collaborators, who reported that free gangliosides (GM1) alter A*β* aggregation, whereby A*β*40 monomers display a preference for binding to GM1-containing hetero-oligomers over GM1-lacking homo-oligomers, while A*β*42 monomers have the opposite preference ^37^. A larger number of free lipids in probing A*β*42-lipid co-assembly and interaction modes has been used in two studies by Urbanc and collaborators ^38,39^. Andrews and collaborators further showed that A*β*42 monomers co-assemble with 12 and 48 free DMPC lipids into stable A*β*-lipid complexes, exhibiting a single A*β*42-DMPC interaction mode of an adhesion to the surface of the DMPC cluster ^38^. Recently, Pandey and collaborators reported that in the process of A*β*42 co-assembly with free lipids, comprised of DMPC, CHOL, and GM1, a lipid bilayer-like structure with a CHOL-rich domain is formed, whereby the only mode of A*β*42 monomer interaction with lipids is an adhesion to the side of the bilayer-like structure, while in addition to the adhesion, A*β*42 dimers get spontaneously inserted into a CHOL-rich domain ^39^. Together, these findings show that MD is an ideal tool for examining early stages of A*β* co-assembly with free lipids and identifying A*β*-lipid interaction modes.

Both *in vitro* and *in silico* studies have shown that in the absence of lipids, A*β*40 and A*β*42 assemble through distinct oligomerization pathways ^5,22^. We posit that the 5% sequence dissimilarity between the two isoform leads not only to their distinct self-assembly pathways, but reflects also in their co-assembly with lipids, which affects both A*β*-A*β* and lipid-lipid association. In this work, we apply ThT fluorescence spectroscopy to examine the effect of the protein to lipid molar concentration (PL) ratio on the kinetics of A*β*40 and A*β*42 fibril formation in a system of pure DMPC lipids and in a binary mixture of DMPC lipids with CHOL. Experimental results demonstrate that the PL ratio of 1:5 inhibits the kinetics of A*β*40 and A*β*42 fibril formation the most. Using the PL ratio of 1:5, we perform extensive explicit-solvent all-atom MD simulations of six initially monomeric A*β*40 and A*β*42 peptides, each co-assembling with 30 free lipids for each of the two lipid systems, DMPC and DMPC/CHOL. To account for the IDP nature of A*β*, five 1 *µ*s-long replica MD trajectories are acquired for each of these four systems, resulting in 20 *×* 1 *µ*s = 20 *µ*s of simulation time. Two control lipids-free systems (with 5 replica MD trajectories per system) and two control A*β*-free systems (with 3 replica MD trajectories per system) are also collected, resulting in combined total simulation time of 36*×* 1 *µ*s = 36 *µ*s. Analysis of all eight systems enables us to derive a comprehensive understanding of how sequence specificity affects A*β*-lipid co-assembly and interaction modes and how these interaction modes are modulated by the presence of CHOL in the DMPC/CHOL system relative to the DMPC system.

## 2 Methods

### 2.1 Experimental Methods

#### Preparation of samples

A*β*40 or A*β*42 (rPeptide) is prepared at a final concentration of 20 *µ*M with 25 *µ*M ThT in 10 mM sodium phosphate buffer (pH 7.4) by diluting a 4 mM stock solution dissolved in DMSO. For experiments in the presence of lipids, homogeneous A*β*-lipid mixtures are prepared using organic solvents. 1,2-dimyristoyl-sn-glycero-3-phosphocholine (DMPC, Avanti Research) and CHOL (Avanti Research) are dissolved in chloroform, while A*β* peptides are dissolved in DMSO. The lipid and A*β* peptide solutions are combined at the desired A*β*:lipid molar ratios, and the organic solvents are removed under a stream of nitrogen (N_2_) gas. Samples are then placed in a lyophilizer overnight to ensure a complete removal of residual solvent. The dried A*β* –lipid films are then hydrated with 10 mM sodium phosphate buffer (pH 7.4) containing 25 *µ*M ThT to prepare samples with final A*β*-to-lipid molar ratios of 1:0.1, 1:1, 1:5, 1:10, and 1:100 for A*β*40:DMPC, A*β*42:DMPC, and A*β*42:(DMPC:CHOL, 7:3).

#### Thioflavin (ThT) fluorescence measurements

ThT fluorescence measurements are performed in Greiner Bio-One 96-well microplates (655900) using a FLUOstar Omega microplate reader (BMG LABTECH) at room temperature, with excitation at 448 nm and emission at 480 nm. Fluorescence readings are collected every 30 min with a gain setting of 1110. Prior to each measurement, plates are orbitally shaken at 700 rpm for 60 s. All systems are measured in duplicate.

### 2.2 Computational Methods

#### Molecular dynamics simulation protocol

Fully atomistic MD simulations in explicit solvent were performed with GROMACS 2021.1^40–46^ using the CHARMM36m force field with its specific TIP3P water model ^47–50^. CHARMM36 force field for lipids was used to model dimyristoylphosphatidylcholine (DMPC), and CHOL ^51–53^. Six A*β*-containing systems were simulated, consisting of six A*β*40 or A*β*42 peptides in the absence of lipids, with 30 DMPC molecules, or with 21 DMPC and 9 CHOL molecules in 150 mM NaCl. Five independent MD trajectories, each 1 *µ*s long, were acquired for each A*β*-containing system. For the lipid-only systems, three independent 1 *µ*s trajectories were acquired for the system containing 30 DMPC molecules, whereas five independent 1 *µ*s trajectories were acquired for the system containing 21 DMPC and 9 CHOL molecules. To maintain consistent sampling between the two lipid-only systems, three trajectories from each system were used in the analyses. The combined simulation time was thus 36 *µ*s. Five distinct A*β*42 conformations were selected from the final 1 ns of previously reported A*β*42 monomer MD trajectories in the absence of lipids ^39^. The corresponding A*β*40 conformations were generated from these A*β*42 structures by removing the two C-terminal residues, I41 and A42. Each resulting conformer was then placed in a (12.5 nm)^3^ simulation box. The N- and C-termini of each A*β*40 and A*β*42 peptide corresponded to NH^+^ and COO^*−*^, respectively. Three Na^+^ ions were added for each A*β*40 and A*β*42 peptide to neutralize the system. The addition of hydrogen atoms to A*β*40 and A*β*42 conformations, the addition of ions to the solvent, and the resolution of atom collisions were performed within GROMACS during the preparation and energy minimization steps. The Verlet cutoff scheme ^54^ and a time step of 2 fs were used during the equilibration and production steps. Energy minimization was performed with the steepest descent method for 100,000 steps and was followed by a 200 ps–long equilibration step at 310 K and 1.0 bar. All production runs used the Nosé-Hoover thermostat ^55,56^ and a Parrinello-Rahman barostat ^57^.

#### A*β* secondary structure

The secondary structure of A*β*40 and A*β*42 in the absence and presence of lipids is analyzed using the STRIDE algorithm ^58^ implemented in the Visual Molecular Dynamics (VMD) software package ^59^. For each of the six systems under study (A*β*40 and A*β*42, each in the absence of lipids, with DMPC lipids, and with DMPC/CHOL lipids), two types of secondary structure analysis are performed: (i) the average coil, strand, turn, and helical content per peptide and (ii) the average per-residue coil, strand, turn, and helical propensities. The helical secondary structure is defined as the sum of three types of helices: *α*-helix (the dominant contribution), *π*-helix, and 3-10 helix. The secondary structure analysis is performed on all A*β*40 and A*β*42 conformations within the time interval of 600-1000 ns of each of the five replica trajectories per system, using frames separated by 1 ns. For each system, the final values are obtained by averaging over the frames of each trajectory, followed by averaging over the five independent trajectories. The error bars represent the standard error of the mean (SEM).

#### A*β* tertiary and quaternary structure

Using VMD ^59^, intra- and intermolecular contact maps of tertiary and quaternary contacts, respectively, were derived to characterize A*β*40 and A*β*42 hexamer structure. A pair of amino acids is considered to be in contact if the distance between the respective C_*α*_ atoms is less than 6 Å. If the two amino acids belong to the same peptide, the contact is recorded as a tertiary contact; otherwise, it is recorded as a quaternary contact. A contact map is thus a two-dimensional matrix of contact probabilities between pairs of residues. A*β*40 and A*β*42 tertiary and quaternary contact maps in the absence of lipids and in the presence of DMPC or DMPC/CHOL were calculated using time frames separated by 1 ns within the time interval of 600-1000 ns of each of the five replica trajectories per system. For each system, contact maps were computed for all relevant frames of each trajectory and then averaged across five replica trajectories. Only time frames in which all six A*β* peptides formed a hexamer were included in the calculation. Of the 2005 frames available across the five replica trajectories per system, 1663, 1296, and 1536 frames were included for A*β*40 in the absence of lipids, in the DMPC system, and in the DMPC/CHOL system, respectively. Similarly, of the 2005 time frames available, only 1894, 1195, and 1312 frames were included for A*β*42 system without lipids, in the DMPC system, and in the DMPC/CHOL system, respectively. A contact map value at (i,j) represents the probability of contact between the amino acids i and j and the standard error of means (SEM) calculated over the frames and trajectories were plotted above and below the main diagonal, respectively. We also computed a contact map difference by subtracting the contact map values at (i,j) obtained in the absence of lipids from the contact map obtained in the presence of lipids for each A*β*40 and A*β*42. In addition, A*β*42 to A*β*40 difference contact maps were calculated by subtracting the A*β*40 contact map from the corresponding A*β*42 contact map. Because A*β*42 contains two additional C-terminal residues, the subtraction was performed over the common 40-residue matrix, with the two additional rows and columns in the resulting A*β*42-sized difference map set to zero.

#### Cluster size distributions

The clustering behavior of the following systems: lipids alone, A*β*40 only, A*β*42 only, A*β*40 with lipids, and A*β*42 with lipids is examined using a 3.5 Å threshold (cutoff) within the GROMACS tool ‘clustsize’, whereby a molecule (A*β*40, A*β*42, DMPC, or CHOL) is identified as part of the same cluster if any of its non-hydrogen atoms are within a distance of 3.5 Å from any of the non-hydrogen atoms of another molecule. The average A*β* oligomer size distributions are calculated using time frames, separated by 1 ns, within 600-1000 ns of each trajectory, followed by an ensemble average over five trajectories per A*β*-containing system and three trajectories per lipid-only system.

#### Per-residue A*β*-lipid contact probabilities

To gain insight into residue-specific interactions between A*β*40 or A*β*42 and lipid molecules, we calculate the average per-residue contact frequencies between A*β*40 or A*β*42 and DMPC or CHOL. Atomic contacts are defined using a distance threshold of 3.5 Å between any non-hydrogen atom of a specific amino acid residue in A*β*40 or A*β*42 and any non-hydrogen atom of DMPC or CHOL. The average number of atomic contacts between a specific residue of A*β*40 or A*β*42 and DMPC or CHOL is calculated using time frames within 600-1000 ns, separated by 1 ns, of all five MD trajectories per system. In the case of A*β* oligomers, the average number of atomic contacts represents an average over the six peptides. The resulting per-residue contact frequencies are obtained by normalizing the average numbers of contacts by the number of lipid molecules in each system, i.e. 30 for DMPC in the DMPC-only system, and 21 for DMPC and 9 for CHOL in the DMPC/CHOL system, to ensure comparability between DMPC and CHOL.

#### Projections of lipid directors on the bicelle normal

The orientational alignment of lipid molecules relative to the normal of the bicelle-like lipid cluster is quantified by projecting the individual lipid molecular axes onto a global director. For each frame, the molecular orientation vectors (**u**_*i*_) for lipids (DMPC and CHOL) are calculated using the ‘compute_directors’ function from MDTraj ^60^, which estimates principal molecular axes based on atomic coordinates. A global bilayer director (**n**) is determined for each frame from the ensemble of lipid molecular axes. Specifically, a second-rank ordering tensor (Q-tensor) is constructed from all lipid orientation vectors, and its principal eigenvector (corresponding to the largest eigenvalue) is taken as the global director. For each lipid molecule in every frame, the scalar projection **u**_*i*_.**n** is computed. This quantity reflects the degree of alignment of each lipid relative to the bilayer normal. The projection values are computed over all lipids, frames, and trajectories within the time window of 900-1000 ns.

#### Density profile of a bicelle

The formation of a lipid bilayer is examined for six systems: (i) 30 DMPC, (ii) 21 DMPC and 9 CHOL, (iii) A*β*40 with 30 DMPC, (iv) A*β*40 with 21 DMPC and 9 CHOL, (v) A*β*42 with 30 DMPC, and (vi) A*β*42 with 21 DMPC and 9 CHOL, whereby all time frames, separated by 1 ns, within 900-1000 ns of replica MD trajectories per system are used in the analysis. For each time frame, a normal to a lipid bilayer, hereby referred to as a director, is determined using the function ‘compute_directors’ from the Python software package MDTraj ^60^. Then, atomic positions of the selected headgroup atoms, i.e. N atoms of DMPC and O atoms of CHOL, are projected onto the bilayer director to obtain a one-dimensional distance distribution, corresponding to the headgroup density profile of the bilayer. We first calculate per-trajectory headgroup density profiles, followed by an ensemble average over all replica MD trajectories.

#### Nematic order parameters of a bicelle

We calculate the nematic order parameter, *S*_2_, using the function ‘compute_nematic_order’ within MDTraj ^60^ to quantify the orientational order of lipids in a bilayer. For each DMPC or CHOL lipid molecule *i* in a lipid assembly of interest, the molecular orientation vector **u**_*i*_ is determined from a molecular inertia tensor defined as

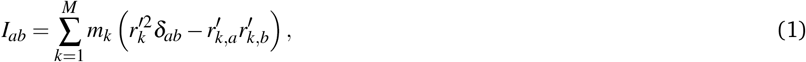

where *m*_*k*_ and 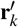 denote the mass and position of each non-hydrogen atom *k* relative to the center of mass of lipid *i*, respectively, and *δ*_*ab*_ is the Kronecker delta. The molecular inertia tensor is then diagonalized, resulting in three eigenvalues and eigenvectors corresponding to the principal moments of inertia and respective axes of rotation. The director of lipid *i*, unit vector **u**_*i*_, is identified as the eigenvector associated with the smallest eigenvalue of *I*_*ab*_, i.e. the smallest rotational moment of inertia around the long molecular axis of the molecule.

A nematic order parameter of the lipid assembly is derived from the alignment tensor, which is calculated using lipid directors **u**_*i*_ as

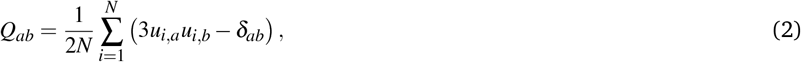

where *N* is the number of lipids in the assembly. The nematic order parameter *S*_2_ is identified as the largest eigenvalue of *Q*, and the corresponding eigenvector is used as a self-consistent global nematic director of the lipid assembly. This formulation is mathematically equivalent to the ensemble average of the second Legendre polynomial, *S*_2_ = ⟨ (3 cos^2^ *θ−* 1)*/*2⟩, where *θ* is the angle between each lipid director and the global nematic director. Because a spontaneously formed lipid assembly may exhibit tilt and/or curvature, the order parameter is computed globally over all lipids without separating leaflets.

Average DMPC and CHOL nematic order parameters are calculated using only DMPC or only CHOL lipid molecules within the assembly, resulting in DMPC and CHOL order parameters, respectively. In addition, the above calculation of tensor *Q* is performed using all DMPC and CHOL lipids in the assembly. The diagonalization of this tensor results in the eigenvector associated with the largest eigenvalue, i.e. a director, which is identified as the normal to the bilayer, unit vector **n**, for all configurations in which lipids form a single cluster, and is used in the calculation of the lipid density profiles and projections of lipid directors onto the bilayer normal.

For each time frame within 900-1000 ns of each MD trajectory of interest, the lipid order parameters are calculated and averaged over all time frames, separated by 1 ns. The final order parameter values are thus obtained as time and ensemble averages over the relevant trajectories per system and the SEM values reflect both time and trajectory-to-trajectory variability.

#### Area per lipid (APL)

For each trajectory frame, the normal of the lipid assembly is determined from the largest eigenvector of the lipid orientational tensor. Lipid headgroup positions, defined by the N-atom of DMPC and the O-atom of CHOL, are then projected onto the normal relative to the center of the lipid assembly and separated into two leaflets according to the sign of the projection. The headgroup coordinates of each leaflet are projected onto the plane perpendicular to the normal of the lipid assembly. The area of each leaflet is calculated by connecting the outermost projected lipid headgroup coordinates to define the leaflet boundary and determining the area enclosed by this boundary using the Convex Hull routine implemented in *SciPy* python library ^61^. The APL value of each leaflet is obtained by dividing the resulting areas by the number of lipids in each leaflet and the mean of the two per-leaflet values is further averaged over 900-1000 ns of each replica trajectory and then across all replica trajectories of each system. The SEM values reflect trajectory-to-trajectory variability of the APL values.

#### Average per-residue solvent accessible surface area (SASA)

For each system with A*β*40 or A*β*42, the average per-residue solvent-accessible surface area (SASA) of A*β* is calculated using the `gmx sasa` tool in GROMACS as a sum of per-residue SASA values divided by the number of residues, which is then averaged over the six peptides in each frame. These average per-residue SASA values are collected for each frame within a time interval of 900-1000 ns of each of the five replica trajectories, where a frame separation of 20 ps is used. Normalized 2D histograms, derived from these SASA values, use Δ*SASA* = 0.00332 nm^2^.

#### A*β*-A*β* and A*β*-lipid potential energies

The A*β*-A*β* and A*β*-lipid potential energies are calculated for each frame within the time interval of 900-1000 ns of each of the five replica trajectories of each A*β*-containing system using frame separation of 20 ps and the `gmx energy` analysis tool in GROMACS. For both potential energy types, the potential energy is calculated as the sum of the short-range Lennard-Jones and Coulomb interaction energies between the corresponding molecular groups. In the case of A*β*-A*β* interactions, the potential energies between all 15 pairs formed by the six A*β* peptides are summed up to obtain the total interpeptide potential energy. In the case of A*β*-lipid interactions, the potential energies between each of the six A*β* peptides and lipid molecules are summed up, including both DMPC and CHOL in the DMPC/CHOL systems. The total potential energy per frame is divided by six to obtain the potential energy per peptide. Normalized 1D and 2D histograms, derived from these potential energy values, use Δ*E* = 5.9 kJ*/*mol.

## 3 Results

In this work, we examine A*β*40 and A*β*42 co-assembly with free lipids using experiment-driven MD simulations. A*β* is set to co-assemble with two systems of free lipids: pure DMPC lipids (the DMPC system) and a binary mixture of DMPC lipids and CHOL at a molar DMPC:CHOL ratio of 7:3 (the DMPC/CHOL system). We posit that the PL ratio is critical for delineating the balance among A*β*-lipid, A*β*-A*β*, and lipid-lipid interactions that lead to specific co-assembly pathways. In the two limiting cases of a very low and a very high PL ratio, we expect the lipid self-assembly into vesicles and A*β* fibril formation, respectively, to proceed mostly unperturbed. In contrast, at intermediate PL ratios around 1:1, we anticipate that A*β*-lipid interactions will affect both A*β* fibril formation and lipid vesicle formation. We first examine A*β*40 and A*β*42 fibril formation kinetics using ThT fluorescence spectroscopy to identify the PL ratio which inhibits fibril formation the most. We then perform MD simulations at thus selected PL ratio to elucidate atomistic details at early stages of A*β*40 and A*β*42 co-assembly with free lipids.

### 3.1 Experimental insights on A*β*40 and A*β*42 fibril formation at different A*β*:lipid molar ratios

Kinetics of A*β*40 and A*β*42 fibril formation in the presence of free lipids is examined by ThT fluorescence spectroscopy (see *Experimental Methods*). In Fig. 1A, normalized ThT fluorescence intensity of A*β*40 fibril formation in the presence of free DMPC lipids versus incubation time, respectively, is displayed at six PL ratios, 1:0, 1:0.1, 1:1, 1:5, 1:10, and 1:100 including the control case with no lipids (black curve). At the lowest PL ratio, A*β*40 fibril formation kinetics is not significantly affected (Fig. 1 A, brown curve). In contrast, at all higher PL ratios, A*β*40 fibril formation is slowed down, and this effect is the strongest at PL ratios of 1:1 and 1:10 (orange and purple curves in Fig. 1 A). Overall, DMPC lipids thus slow down A*β*40 fibril formation by increasing the lag time, possibly due to A*β*40 interactions with DMPC lipids at early stages of co-assembly.

**Figure 1.**
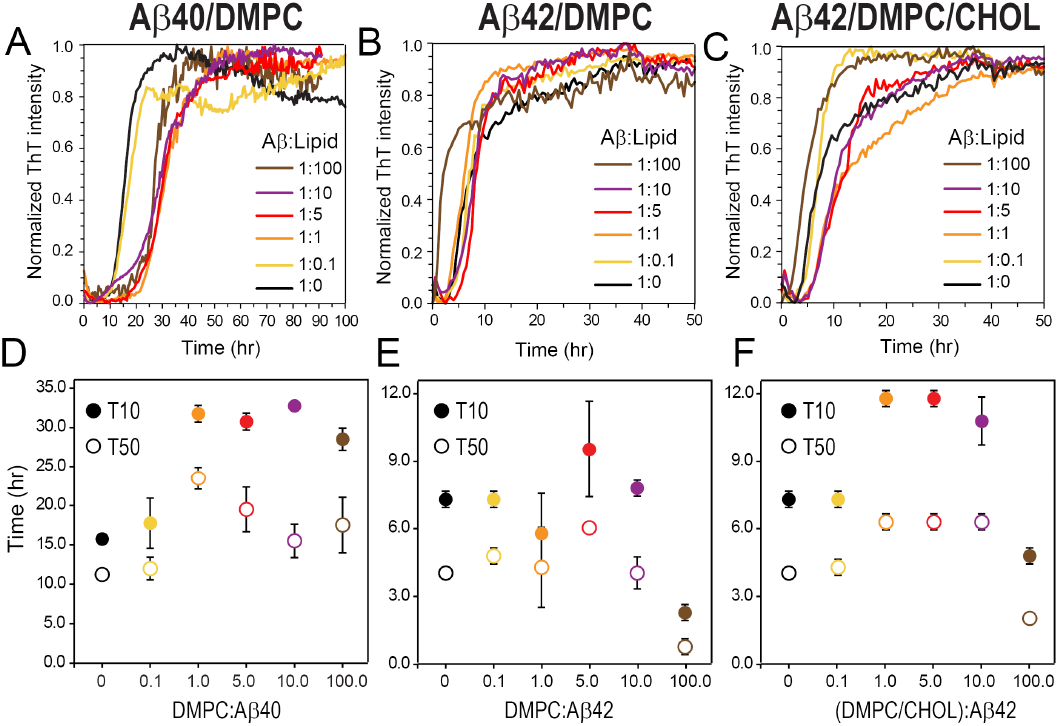
Aggregation kinetics of 20 *µ*M A*β* at different A*β*:lipid ratios. ThT fluorescence intensity of (A) A*β*40 and (B) A*β*42 incubated in the absence and presence of DMPC lipids in dependence on incubation time. (C) ThT fluorescence intensity of A*β*42 incubated with a mixture of DMPC–CHOL lipids at the molar DMPC:CHOL ratio of 7:3 in dependence on incubation time. Molar A*β*:lipid concentration ratios of 1:0 are shown as black, 1:0.1 as yellow, 1:1 as orange, 1:5 as red, 1:10 as purple, and 1:100 as brown curves. (D-F) Quantitative analysis of A*β*40 and A*β*42 aggregation kinetics in the presence of DMPC and DMPC/CHOL mixtures shown in (A-C). T10 (open circles) and T50 (solid circles), representing the times required to reach 10% and 50% of maximal ThT fluorescence intensity, respectively, are shown for (C) A*β*40 with DMPC, (D) A*β*42 with DMPC, and (E) A*β*42 with DMPC/CHOL mixtures (DMPC:CHOL = 7:3) at A*β*:lipid ratios of 1:0 (black), 1:0.1 (yellow), 1:1 (orange), 1:5 (red), 1:10 (purple), and 1:100 (brown). Data represent the mean *±* standard deviation (*n* = 2).

In the absence of lipids, A*β*42 is known to form fibrils faster than A*β*40. Fig. 1B shows normalized ThT fluorescence intensity of A*β*42 fibril formation in the presence of free DMPC lipids versus incubation time. Again, six PL ratios, 1:0, 1:0.1, 1:1, 1:5, 1:10, and 1:100, are probed, including the control case with no lipids (black curve). Fig. 1B demonstrates that at the lowest PL ratio of 1:100 (brown curve), the speed of A*β*42 fibril formation is significantly increased relative to the control (black curve). Thus, in this lipid-dominant regime, the rate of A*β*42 fibril formation is increased unlike in the case of A*β*40. Interestingly, at higher PL ratios, A*β*42 fibril formation rates are not strongly affected by DMPC lipids. Only at the PL ratio of 1:5 (red curve), the kinetics of A*β*42 fibril formation is somewhat slower than in the control case (black curve). These findings show that A*β*-lipid interactions and co-assembly pathways are isoform specific, i.e. A*β*40 and A*β*42 interact with DMPC lipids in distinct ways.

A recent MD study by Pandey and collaborators reported that A*β*42 dimers show a preference for interacting with CHOL over DMPC lipids ^39^. We wondered whether A*β*42 fibril formation is affected by lipids to a greater extent in the DMPC/CHOL than in the DMPC system. Fig. 1 displays ThT fluorescence intensity of A*β*42 fibril formation in the presence of DMPC lipids versus incubation time, respectively, at six PL ratios, 1:0, 1:0.1, 1:1, 1:5, 1:10, and 1:100, including the control case with no lipids (black curve). Akin to the DMPC system, Fig. 1C demonstrates that at the lowest PL ratio of 1:100 (brown curve), the speed of A*β*42 fibril formation in the DMPC/CHOL system is increased relative to the control case (black curve). At the PL ratio of 1:0.1, the DMPC lipids mixed with CHOL do not affect the rate of A*β*42 fibril formation. In contrast, the rate of A*β*42 fibril formation is clearly decreased at PL ratios of 1:1, 1:5, and 1:10, demonstrating that the presence of CHOL in the DMPC/CHOL system results in a decreased rate of A*β*42 fibril formation, consistent with the observed affinity of A*β*42 for CHOL.

A more quantitative analysis shows that for A*β*40, both times to reach 10% and 50% of the maximum ThT fluorescence intensity significantly increase in the presence of DMPC lipids (Fig. 1C). For A*β*42, in contrast, no significant increase is observed except at the PL ratio of 1:5 (Fig. 1D). At the PL ratio of 1:100, A*β*42 fibril formation is significantly faster than in the absence of DMPC lipids (Fig. 1D, brown versus black circles). Importantly, when CHOL is present, a mixture of DMPC lipids with CHOL significantly inhibits A*β*42 fibril formation at A*β*42:lipid ratios of 1:1, 1:5, and 1:10 (Fig. 1F, orange, red, and purple circles). Interestingly, while at the PL ratio of 1:100, A*β*42 fibril formation is faster than in the absence of lipids (Fig. 1F, brown versus black circles), the increase in A*β*42 fibril formation rate is smaller than in the presence of DMPC without CHOL (Fig. 1E, brown versus black circles).

The above experimental data on the kinetics of A*β*40 and A*β*42 fibril formation demonstrate that the PL ratio of 1:5 affects fibril formation of A*β*40 in the presence of DMPC and of A*β*42 in the presence of DMPC/CHOL mixtures. Effects of DMPC and DMPC/CHOL lipid systems on A*β*40 and A*β*42 fibril formation kinetics at the PL ratio of 1:5 are shown in Fig. 2 in more detail. Clearly, pure DMPC lipids slow down A*β*40 fibril formation (Fig. 2A, blue curves), however, this effect is much stronger for DMPC/CHOL mixtures (Fig. 2A, green curves). In contrast, the effect of lipids on A*β*42 fibril formation is much smaller (Fig. 2B). Pure DMPC lipids do not affect the rate of A*β*42 fibril formation, although the saturated ThT intensity is increased (Fig. 2B, orange curves). In the DMPC/CHOL system, A*β*42 fibril formation is slower (Fig. 2B, purple curves), but the effect of lipids on A*β*42 is much weaker than the effect on A*β*40 fibril formation rate.

**Figure 2.**
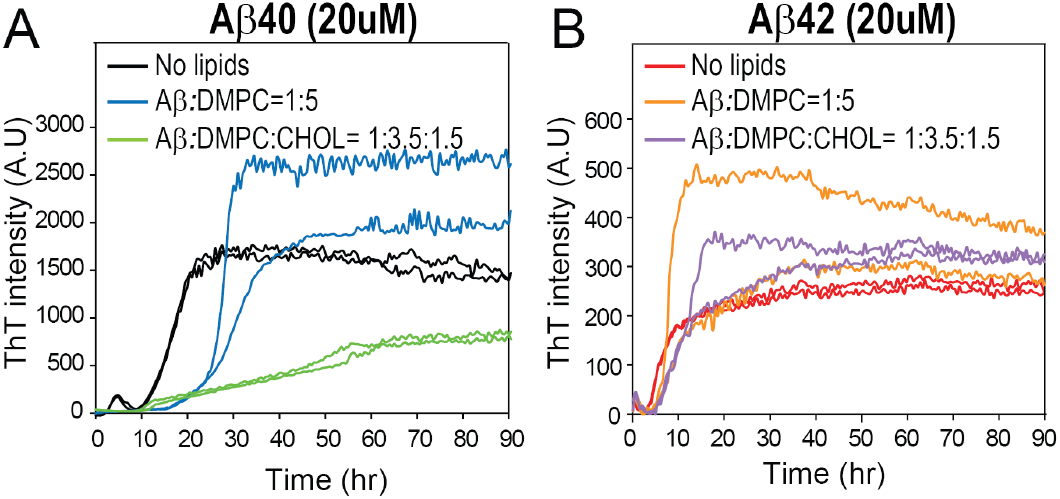
Effects of lipids on aggregation kinetics of 20 *µ*M A*β*. ThT fluorescence intensity in dependence on incubation time of (A) A*β*40 and (B) A*β*42 alone (black and red curves), in the presence of DMPC lipids (blue and orange curves), and in the presence of DMPC-CHOL lipid mixture at the DMPC:CHOL molar ratio of 7:3 (green and purple curves). Data for the two repeated measurements per system are included.

While ThT fluorescence spectroscopy probes A*β* fibril formation in the absence and presence of lipids on timescales of hours, it cannot elucidate processes that occur at early stages of A*β*-lipid co-assembly, which lead to these experimentally observed distinct effects of lipids on A*β* fibril formation. We here posit that the effect of DMPC lipids and DMPC/CHOL mixtures on A*β*40 versus A*β*42 fibril formation arise as a consequence of isoform-specific interactions with DMPC and CHOL molecules at early stages of co-assembly, which can be examined through MD simulations.

### 3.2 MD insights into A*β* co-assembly with lipids

We examine the early stages of co-assembly of A*β*40 and A*β*42 with free lipids by performing MD simulations of eight systems: lipids-only systems containing (i) 30 DMPC lipids or (ii) 21 DMPC and 9 CHOL lipids; A*β*40 systems containing six peptides (iii) without lipids, (iv) with 30 DMPC lipids, or (v) with 21 DMPC and 9 CHOL lipids; and A*β*42 systems containing six peptides (vi) without lipids, (vii) with 30 DMPC lipids, or (viii) with 21 DMPC and 9 CHOL lipids. Lipids-only systems (i) and (ii), and systems without lipids (iii) and (vi) are control systems that allow for studying the effect of A*β* on lipid assembly and vice versa. As explained in *Methods*, we acquired 3 replica MD trajectories for system (i) and 5 replica MD trajectories for systems (ii)-(viii) to feed the statistical analysis. For each of the eight systems under investigation, movies of the assembly dynamics for all replica MD trajectories are included as *Supplementary Information*.

In the analysis of MD trajectories below, we will use the following abbreviations for A*β* regions: NTR for the N-terminal region D1-D7, CHC for the central hydrophobic cluster L17-A21, MHR for the mid-hydrophobic region I31-M35, and CTR for the C-terminal region V39-V40 in A*β*40 or V39-A42 in A*β*42.

#### 3.2.1 Lipids affect A*β* hexamer stability in isoform-specific way

We monitor the time evolution of A*β* oligomer size distributions in the absence and presence of lipids to examine how long it takes for a steady state to emerge (Figures S1, S2, and S3). In all systems, A*β*40 and A*β*42 assemble into oligomers of different sizes within the first several hundred nanoseconds. The early stages of oligomerization are characterized by substantial trajectory-to-trajectory variability, indicating that A*β* assembly follows multiple stochastic pathways. A*β*40 and A*β*42 form larger oligomers over time, with hexamers becoming the dominant species by ~600 ns of simulations both in the absence and presence of lipids. The oligomer size distributions appear to be in a quasi-steady state during 600-1000 ns. Thus, all subsequent A*β* structural analyses are performed using the time interval of 600-1000 ns of all replica MD trajectories of each A*β*-including system.

We then examine self-assembly of lipids in the DMPC and DMPC/CHOL systems in the absence of A*β* by monitoring time evolution of the lipid cluster size distribution (Figure S4). In both lipid-only systems, lipids rapidly assemble into a single cluster by 200 ns in each of the three replica trajectories per system, which remains stable for the remainder of the simulations.

Time evolution of the total cluster size distribution for the systems of A*β* with lipids in Figure S5 allows us to identify the time window, in which A*β* and lipids co-assemble to form a single complex. In all four systems of A*β* with lipids, the total cluster size distributions evolve from multiple smaller clusters at early simulation times toward a single cluster. A*β*40 co-assembling with DMPC lipids converge into a single cluster by 400 ns of simulations in all five replica trajectories. The CHOL-containing lipid mixture further accelerates A*β*40-lipid co-assembly, whereby all trajectories form a single cluster by 200 ns. In contrast, A*β*42 co-assembly with lipids proceeds more slowly than A*β*40-lipid co-assembly. A*β*42 co-assembly with pure DMPC lipids is the slowest; a single cluster forms by 900 ns. In the DMPC/CHOL system, a single A*β*42-lipid cluster emerges by 700 ns. These results indicate that A*β*40 promotes faster co-assembly with lipids than A*β*42. By 900 ns of simulations, a single A*β*-lipid complex emerges in all replica MD trajectories of all four systems. Therefore, time interval 900-1000 ns is used in the analysis of lipid assemblies both in the absence and presence of A*β*.

To determine how free lipids influence early A*β* self-assembly, we calculate A*β*40 and A*β*42 oligomer size distributions in the absence and presence of lipids (Figure 3, panels A and B). For both A*β*40 and A*β*42, hexamer represents the dominant species whereas the populations of monomers and smaller oligomers remain low in all six A*β*-containing systems. A*β*40 co-assembly with lipids in the DMPC/CHOL system produces the highest hexamer propensity whereas A*β*40 co-assembly both in the absence of lipids and with pure DMPC lipids results in somewhat lower hexamer populations that are comparable to each other (Figure 3, panel A). A*β*42 exhibits the highest hexamer propensity in the absence of lipids. In the DMPC system, A*β*42 hexamer population is reduced at the expense of increased monomer and trimer populations, whereas in CHOL-containing lipid mixture, A*β*42 hexamer population increases alongside the occurrence of minor dimer and tetramer populations. Thus, both A*β*40 and A*β*42 assemble into stable hexamers, whereby lipids modulate hexamer stability in an isoform- and composition-dependent way. A*β*40 and A*β*42 oligomer size distributions are compared at three different conditions in Figure 3 panels C, D, and E. In the absence of lipids, A*β*42 has a higher hexamer population than A*β*40. Both isoforms have a small trimer population, but only A*β*40 is characterized by a non-zero monomer population (Figure 3, panel C). Figure 3, panels D and E shows that in the presence of lipids, A*β*40 exhibits a higher hexamer population than A*β*42. Thus, lipids modulate A*β* oligomer size distributions in an isoform- and lipid composition-dependent manner.

**Figure 3.**
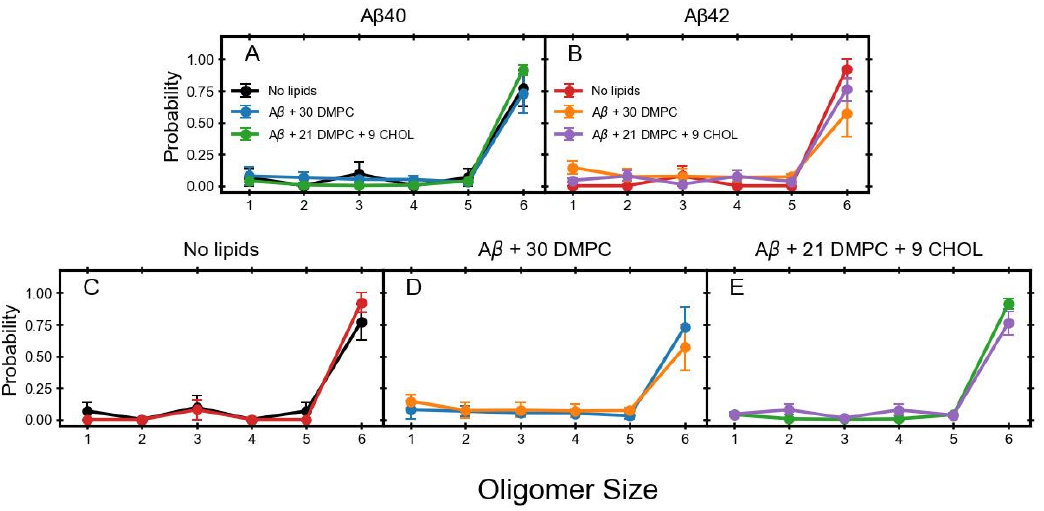
(A,B) The effect of lipids on the oligomer size distribution of (A) A*β*40 and (B) A*β*42. (C-E) A*β*40 vs A*β*42 oligomer size distribution (C) in the absence of lipids, in the presence of (D) DMPC, (E) DMPC/CHOL lipids. The final values are averaged over all the trajectories and selected frames, with error bars indicating the standard error of mean (SEM)

#### 3.2.2 Lipids affect secondary structure in A*β*40 more than in A*β*42

To elucidate the effects of lipids on the secondary structure of A*β*40 and A*β*42, we calculate the secondary structure propensities of both A*β* isoforms in the absence of lipids and in the DMPC and DMPC/CHOL systems. The secondary structure, i.e. coil, *β*-strand, turn, and helix, is analyzed in two ways, by calculating the average secondary structure content per peptide and average per-residue secondary structure propensities. The effect of DMPC and DMPC/CHOL lipids on the average secondary structure content of A*β*40 and A*β*42 is shown in Figure 4 (panels A and B). In A*β*40, the average coil content decreases in the presence of DMPC lipids and even more so in the presence of DMPC/CHOL lipid mixture (Figure 4, panel A). This decrease is accompanied by an increase in *β*-strand, turn, and helical content. The average *β*-strand in A*β*40 is the highest in the presence of DMPC lipids whereas the average turn and helical propensities are the highest in the presence of DMPC/CHOL lipid mixture. In contrast, the average secondary structure content of A*β*42 is less sensitive to lipids (Figure 4, panel B). The average coil and *β*-strand contents in A*β*42 remain comparable across the three conditions, with a minor decrease in the CHOL-containing lipid mixture. The average turn propensity in A*β*42 slightly increases in the DMPC/CHOL system, whereas the average helical content remains low across all three conditions and is nearly absent in the DMPC system. Thus, lipids reduce the structural disorder of A*β*40 but do not significantly affect the average secondary structure content of A*β*42.

**Figure 4.**
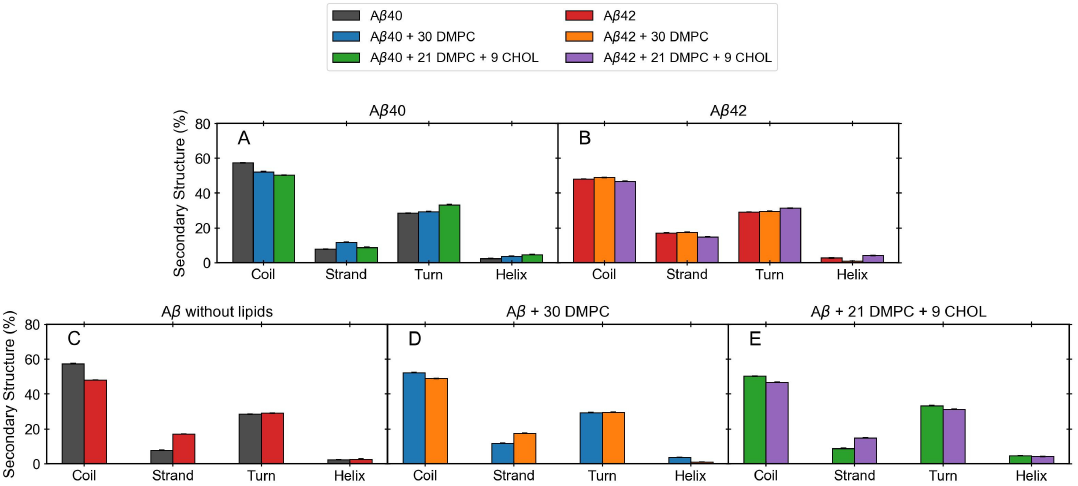
(A,B) The effect of lipids on the secondary structure propensities of (A) A*β*40 and (B) A*β*42. (C-E) Secondary structure propensities of A*β*40 vs A*β*42 (C) in the absence of lipids, in the presence of (D) DMPC, (E) DMPC/CHOL system. The final values are averaged over all the trajectories and selected frames, with error bars indicating the standard error of mean (SEM).

The average secondary structure content of A*β*40 and A*β*42 is compared in Figure 4C-E. Within the time interval of the analysis, 600-1000 ns, the assembly state of A*β*40 and A*β*42 is dominated by hexamers. In the absence of lipids, A*β*40 has a higher average coil content than A*β*42 and A*β*42 has a substantially higher average *β*-strand content than A*β*40 (Figure 4C). The average turn content is comparable between the two isoforms, and the helical content is low for both A*β*40 and A*β*42. Thus, in the absence of lipids, A*β*42 hexamers are more structured than A*β*40 hexamers, primarily due to the enhanced *β*-strand content. The presence of lipids does not change the above qualitative findings, however, the secondary structure content in A*β*40 and A*β*42 hexamers becomes more comparable than in the absence of lipids (Figure 4D-E). The effects of pure DMPC lipids and a mixture of DMPC/CHOL lipids on the average secondary structure content are comparable, however, the presence of CHOL results in somewhat higher, albeit small, average helical content in both A*β*40 and A*β*42 hexamers.

Figure S6A shows the effects of lipids on average per-residue secondary structure propensities. In A*β*40, the presence of lipids reduces coil propensities, most prominently within the CHC (particularly in region F19-A21), and increases *β*-strand propensities, in particular in the DMPC system (Figure S6A, left column). A*β*40 region Y10-E11 also exhibits higher *β*-strand propensities in the presence of lipids, most notably in the DMPC/CHOL system. In addition to *β*-strand propensities, lipids also enhance A*β*40 turn propensities. In both lipid systems, region E22-S26 exhibits increased turn propensities. In the DMPC/CHOL system, turn propensities increase also in region H14-Q15. A*β*40 helical propensities remain low but increase locally in region E22-N27, particularly in the presence of CHOL.

In A*β*42, lipid-induced changes are concentrated primarily within the MHR, region V36-G38, and residues V39-V40 (Figure S6A, right column). Overall, coil propensities remain comparable across the three conditions. Nonetheless, DMPC lipids reduce coil propensities in region V36-V40 and increase *β*-strand propensities in regions L34-M35 and V39-V40. In the DMPC/CHOL system, A*β*42 *β*-strand propensities are decreased in regions G33-M35 and V39-V40 while in region F4-H6, *β*-strand propensities increase. Lipids also alter turn propensities in A*β*42 in a lipid composition-dependent manner. DMPC lipids increase turn propensities in region V36-G38, whereas in the presence of CHOL turn propensities in this same region decrease and increase in region F19-A21. Helical propensities remain low along the entire A*β*42 sequence in all three conditions. In the absence of lipids, non-zero helical propensities are observed in region G33-M35. These propensities are nearly abolished in the DMPC system but are enhanced in the DMPC/CHOL system, which also increases helical propensities in region A30-I32.

Figure S6B compares per-residue secondary-structure propensities of A*β*40 and A*β*42 in three different environments. In the absence of lipids (Figure S6B, leftmost column), A*β*40 and A*β*42 exhibit comparable coil propensities in region D1-Q15. The most prominent isoform-dependent differences occur in regions V18-A21 and D23-V40, where A*β*42 exhibits lower coil propensities than A*β*40, indicating reduced disorder along the central and C-terminal parts of A*β*42. A*β*42 also exhibits higher *β*-strand propensities than A*β*40 in regions R5-D7, K16-M35, and V39-V40, with the most extensive increase spanning region K16-M35. In contrast to these sweeping differences in coil and *β*-strand propensities, differences in turn propensities between the two isoforms are more localized. A*β*40 exhibits higher turn propensities than A*β*42 in regions H6-G9 and L17-F20, whereas A*β*42 exhibits higher turn propensities than A*β*40 in regions D23-S26 and M35-V39. Helical propensities remain very low in both isoforms, whereby A*β*40 shows higher propensities than A*β*42 in region E22-S26 and A*β*42 displays higher propensities than A*β*40 in region G33-V36. Thus, in the absence of lipids, A*β*42 is more ordered than A*β*40, primarily due to the overall decrease in coil propensities and enhancement of *β*-strand propensities in the broad region K16-V40.

In the DMPC system (Figure S6B, central column), the secondary structure propensities of A*β*40 and A*β*42 are more comparable to each other than in the absence of lipids. Per-residue coil propensities are similar along most of the sequence, although A*β*40 retains higher coil propensities than A*β*42 in regions K16-V18 and M35-V40. A*β*42 exhibits higher *β*-strand propensities than A*β*40 in regions K16-F19, L34-M35, and V39-V40. Turn propensities are also comparable between the two isoforms, although A*β*40 exhibits higher turn propensities than A*β*42 in region L17-F20 and A*β*42 shows higher turn propensities than A*β*40 in region V36-G38. The most distinct features between the two isoforms are observed in per-residue helical propensities. Helical propensities are nearly absent in A*β*42, whereas A*β*40 retains non-zero helical propensities in regions E22-S26 and G33-V36. While DMPC lipids reduce the overall differences in the secondary structures propensities between A*β*40 and A*β*42, the distribution of secondary structure propensities along the sequence remains isoform specific.

In the DMPC/CHOL system (Figure S6B, rightmost column), A*β*40 exhibits higher per-residue coil propensities than A*β*42 in regions K16-V18 and A30-V40. A*β*42 is characterized by higher *β*-strand propensities than A*β*40 in regions F4-D7, Q15-F19, A30-I32, at M35, and in region V39-V40. In region Y10-E11, A*β*40 exhibits higher *β*-strand propensities than A*β*42. Overall, both isoforms are characterized by comparable per-residue turn propensities. Whereas A*β*40 exhibits somewhat higher turn propensities than A*β*42 in regions A2-S8 and L17-V18, A*β*42 shows slightly higher turn propensities than A*β*40 in the CTR. Helical propensities remain low in both isoforms but differ in their distribution along the sequence. A*β*40 shows higher helical propensities than A*β*42 in region L17-S26, whereas A*β*42 displays higher helical propensities than A*β*40 in region G33-G37. Thus, in the DMPC/CHOL system, A*β*40 possesses more disordered secondary structure with higher coil propensities than A*β*42. In comparison, A*β*42 exhibits higher *β*-strand propensities than A*β*40. While helical propensities remain low, their distribution along the sequence is isoform specific with A*β*40 and A*β*42 showing non-zero helical propensities in the CHC and MHR, respectively.

The above results demonstrate that lipids affect the secondary structure of A*β*40 more than the secondary structure of A*β*42. The presence of CHOL in the DMPC/CHOL system further modifies lipid-induced secondary structure changes by promoting turn and helical content at the expense of *β*-strand in an isoform-specific manner.

#### 3.2.3 Lipids affect the tertiary structure of A*β* hexamers in a lipid system- and isoform-specific manner

Intramolecular contact maps of A*β*40 and A*β*42 hexamers in the absence of lipids, in the DMPC system, and in the DMPC/CHOL system are shown in Figure 5, with a detailed description in *Supplementary Results*. The analysis of A*β*40 and A*β*42 intramolecular contact maps demonstrates that in all three environments, tertiary contacts form with a rather low probability that does not exceed 0.1, consistent with the IDP nature of these peptides. Key observations are summarized below.

**Figure 5.**
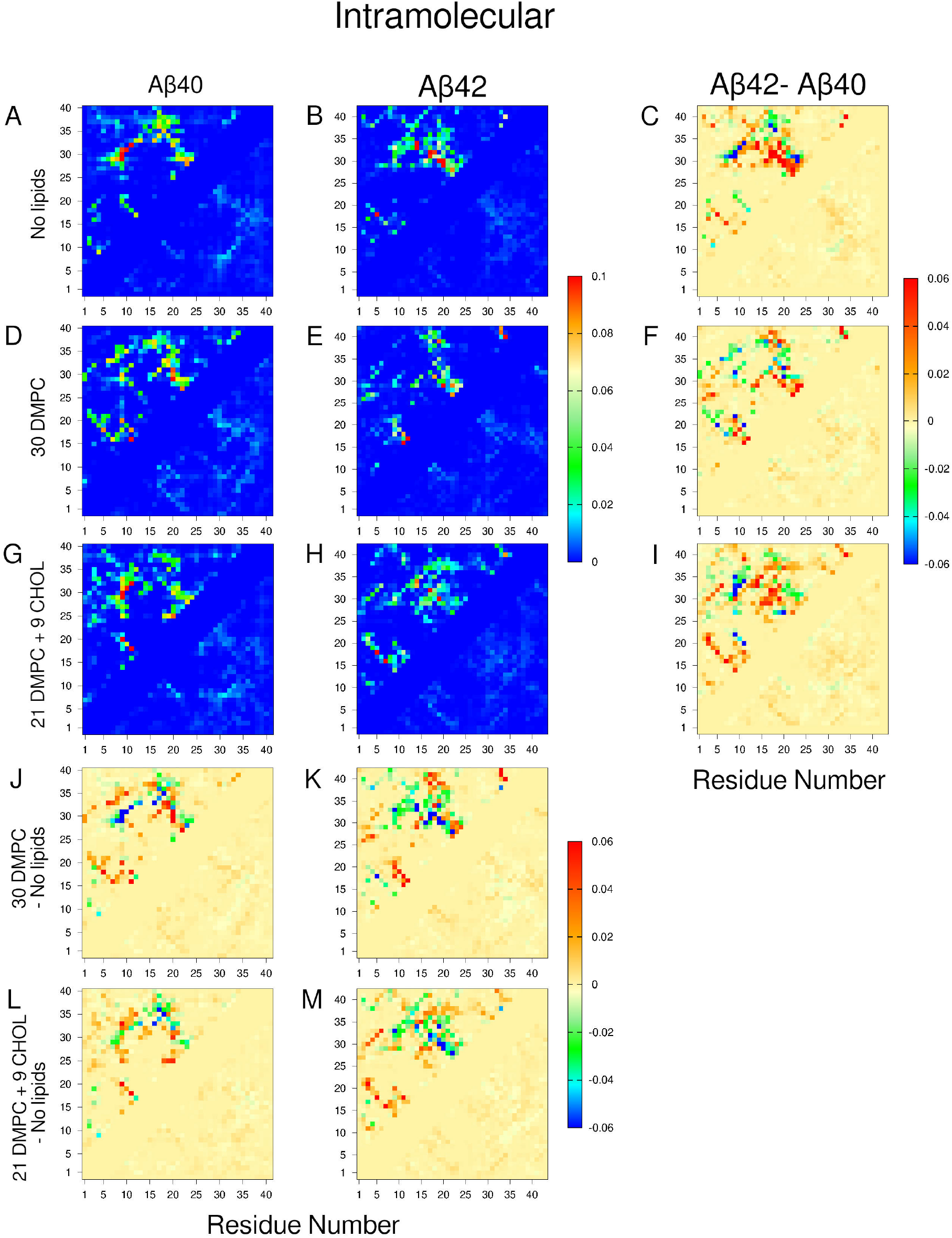
The effect of lipids on tertiary structure of (A, D, G) A*β*40 and (B, E, H) A*β*42 under three conditions (A,B) No lipids, (D,E) with DMPC lipids, and (G,H) with DMPC/CHOL lipids. (J-M) show lipids minus no lipids contact maps to elucidate the tertiary structure changes in presence and absence of lipids. (C,F,I) show the difference A*β*42 - A*β*40 in different lipid environments. The color scale shows the contact probabilities. The final values are calculated over the selected frames and trajectories with error bars representing the standard error of mean (SEM)

In the lipid-free environment, the dominant tertiary contacts in both A*β*40 and A*β*42 hexamers are centered within the central folding region, A21-A30, which results in proximate intramolecular contacts between the CHC and MHR. These CHC-MHR contacts are substantially more probable in A*β*42 than in A*β*40 hexamers, as evident from the A*β*42 to A*β*40 difference contact map in Figure 5A-C. The only common lipid-induced feature in A*β*40 and A*β*42 hexamers is an enhanced participation of the NTR in tertiary contact formation.

Pure DMPC lipids enhance tertiary contacts between the CHC and MHR and promote a distinct turn-like structure between the A2-D7 region and the CHC in A*β*40 hexamers, as shown in Figure 5(D,J). This NTR-CHC turn-like structure is absent from A*β*42 hexamers co-assembled with pure DMPC lipids, as can be noted in the A*β*42 to A*β*40 difference contact map in Figure 5F. Unlike in A*β*40 hexamers, pure DMPC lipids inhibit tertiary contacts between the CHC and MHR in A*β*42 hexamers.

In the DMPC/CHOL system, intramolecular contact map of A*β*40 hexamers resembles more closely the intramolecular contact map of A*β*40 hexamers formed in the lipid-free environment than the corresponding intramolecular contact map in pure DMPC lipids (Figure 5A,D,G). In contrast, A*β*42 hexamers co-assembled with DMPC lipids and CHOL, the CHC-MHR contacts are further reduced relative to the environment of pure DMPC lipids (Figure 5E,H).

#### 3.2.4 Lipids reshape the quaternary structure of A*β* hexamers in a lipid system- and isoform-specific manner

Intermolecular contact maps of A*β*40 and A*β*42 hexamers in the absence of lipids, in the DMPC system, and in the DMPC/CHOL system are shown in Figure 6. A detailed description of the effect of lipids on the quaternary structure of A*β*40 and A*β*42 hexamers can be found in *Supplementary Results*. Similar to intramolecular contact maps, intermolecular contact maps indicate that quaternary contacts occur with a low probability that does not exceed 0.1. In both lipid systems, quaternary contacts of A*β*40 and A*β*42 hexamers are reduced, indicating that A*β*-lipid interactions are competing with A*β*-A*β* interactions. Just like in the case of the tertiary structure, lipid-induced changes in the quaternary structure of A*β* hexamers are isoform- and lipid system-specific. A summary of key observations is given below.

**Figure 6.**
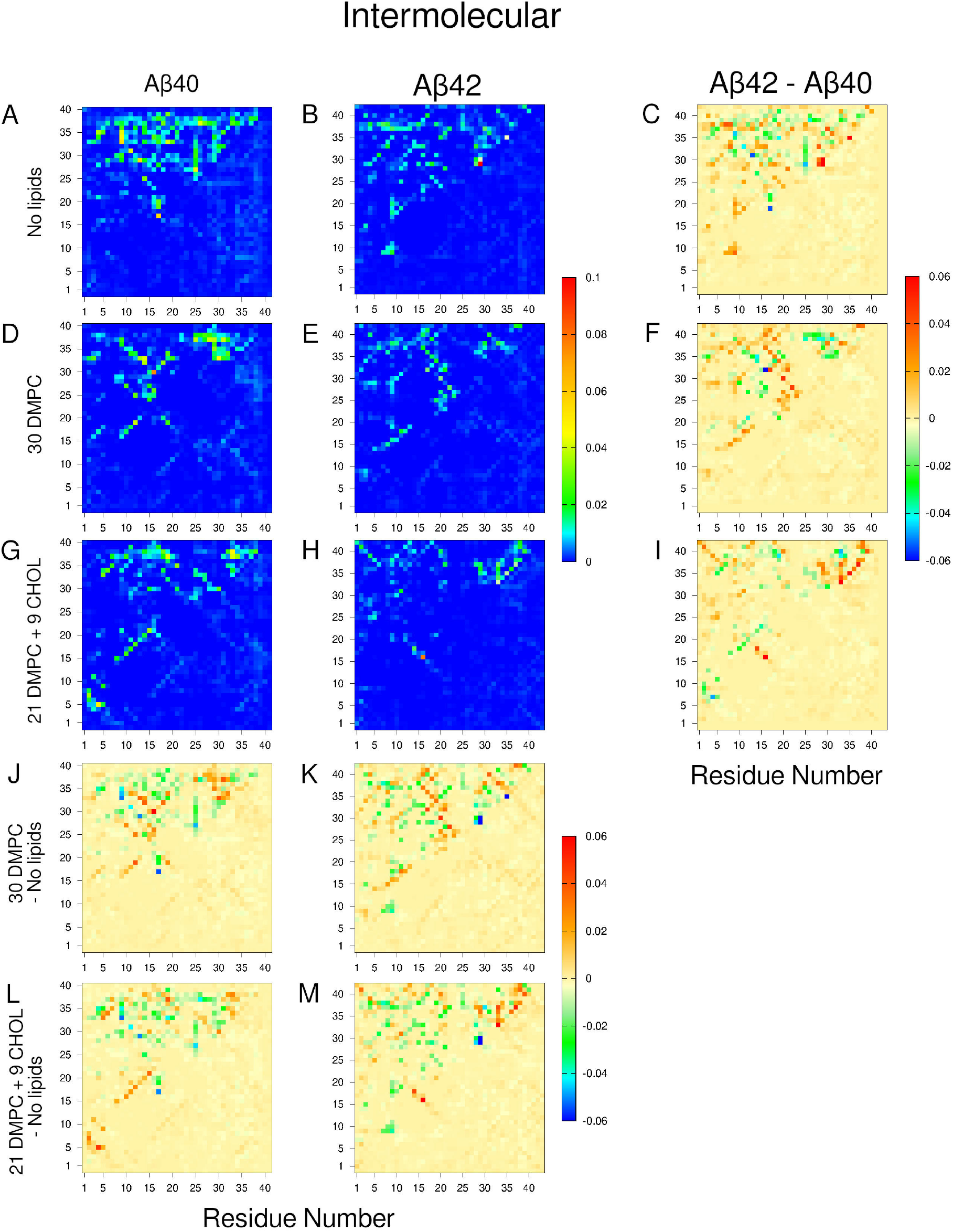
The effect of lipids on quaternary structure of (A, D, G) A*β*40 and (B, E, H) A*β*42 under three conditions (A,B) No lipids, (D,E) with DMPC lipids, and (G,H) with DMPC/CHOL lipids. (J-M) show lipids minus no lipids contact maps to elucidate the quaternary structure changes in presence and absence of lipids. (C,F,I) show the difference A*β*42 - A*β*40 in different lipid environments. The color scale shows the contact probabilities. The final values are calculated over the selected frames and trajectories with error bars representing the standard error of mean (SEM)

In the lipid-free environment, the most prominent quaternary contacts in A*β*40 hexamers are centered around contact L17-L17, bringing pairs of the CHC regions into proximity. Quaternary contacts in A*β*40 hexamers are also characterized by a broad distribution of contacts along the sequence, including contacts between regions G25-G29 and G33-V39 and contacts between the NTR and the CHC. In A*β*42 hexamers, the most prominent quaternary contacts are centered around G29-G29 and their distribution along the sequence shows a predominance of parallel, in-register quaternary contacts, indicating that unlike A*β*40, A*β*42 peptides form parallel intermolecular strands. This observation is supported by the A*β*42 to A*β*40 difference contact map in Figure 6C.

Pure DMPC lipids significantly reduce the number of quaternary contacts in A*β*40 hexamers and to a lesser extent also in A*β*42 hexamers (Figure 6D-F,J,K). The A*β*42 to A*β*40 difference contact map reveals distinct residue-specific contact preferences of hexamers formed by the two isoforms (Figure 6F). A*β*40 hexamers are stabilized by quaternary contacts between regions H14-K16 and K28-I31 and additional quaternary contacts between the MHR and the CTR. On the other hand, A*β*42 hexamer formation is driven by intermolecular contacts between the CHC and region E22-A32, in addition to the G37-A42 contact.

In the DMPC/CHOL system, A*β*40 hexamers are characterized by broadly distributed intermolecular contacts whereas A*β*42 hexamers favor more specific, parallel in-register intermolecular contacts along the MHR and CTR that are not present in A*β*40 hexamers (Figure 6G-I,L,M).

#### 3.2.5 Lipids self-assemble into a layered micelle (bicelle)

Figure S4 shows that in both lipid-only systems, lipids self-assemble into a single lipid cluster within the first 200 ns of simulations in all three replica trajectories per system, whereas in the presence of A*β*40 or A*β*42, A*β* co-assembly with lipids into a single cluster requires significantly longer simulation time, up to 900 ns (Figure S5). Visual inspection reveals lipid clusters resembling micelles with a layered structure, which we refer to hereafter as bicelles. We analyzed the morphology of bicelles and the results of this analysis are reported in Figure 7A-V and Table 1.

**Table 1.** Lipid alignment, bilayer thickness *D*_*B*_, and area per lipid (APL) for lipid-only and A*β*–lipid systems. Alignment is reported as the fraction of lipids satisfying 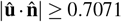. Thickness and APL values are reported as mean *±* SEM across trajectories over 900-1000 ns.

| System | Aligned | $D_B$-DMPC (nm) | $D_B$-CHOL (nm) | APL (nm<sup>2</sup>) |
| --- | --- | --- | --- | --- |
| DMPC | 0.453 | 3.65 ± 0.07 | | 0.8076 ± 0.0011 |
| +6 Aβ40 | 0.548 | 3.58 ± 0.22 | | 0.9317 ± 0.0366 |
| +6 Aβ42 | 0.520 | 3.57 ± 0.25 | | 0.8967 ± 0.0439 |
| DMPC/CHOL | 0.526 | 3.45 ± 0.03 | 1.94 ± 0.26 | 0.6760 ± 0.0011 |
| +6 Aβ40 | 0.604 | 3.66 ± 0.12 | 2.67 ± 0.37 | 0.7262 ± 0.0401 |
| +6 Aβ42 | 0.562 | 3.42 ± 0.18 | 2.67 ± 0.41 | 0.7662 ± 0.0407 |
| DMPC/CHOL/GM1<sup>1</sup> | 0.682 | 4.27 ± 0.018 | 3.17 ± 0.17 | 0.5233 ± 0.0014 |
| +1 Aβ42 | 0.703 | 4.28 ± 0.016 | 3.15 ± 0.13 | 0.5372 ± 0.0019 |
| +2 Aβ42 | 0.682 | 4.18 ± 0.11 | 3.15 ± 0.14 | 0.5474 ± 0.0094 |

**Figure 7.**
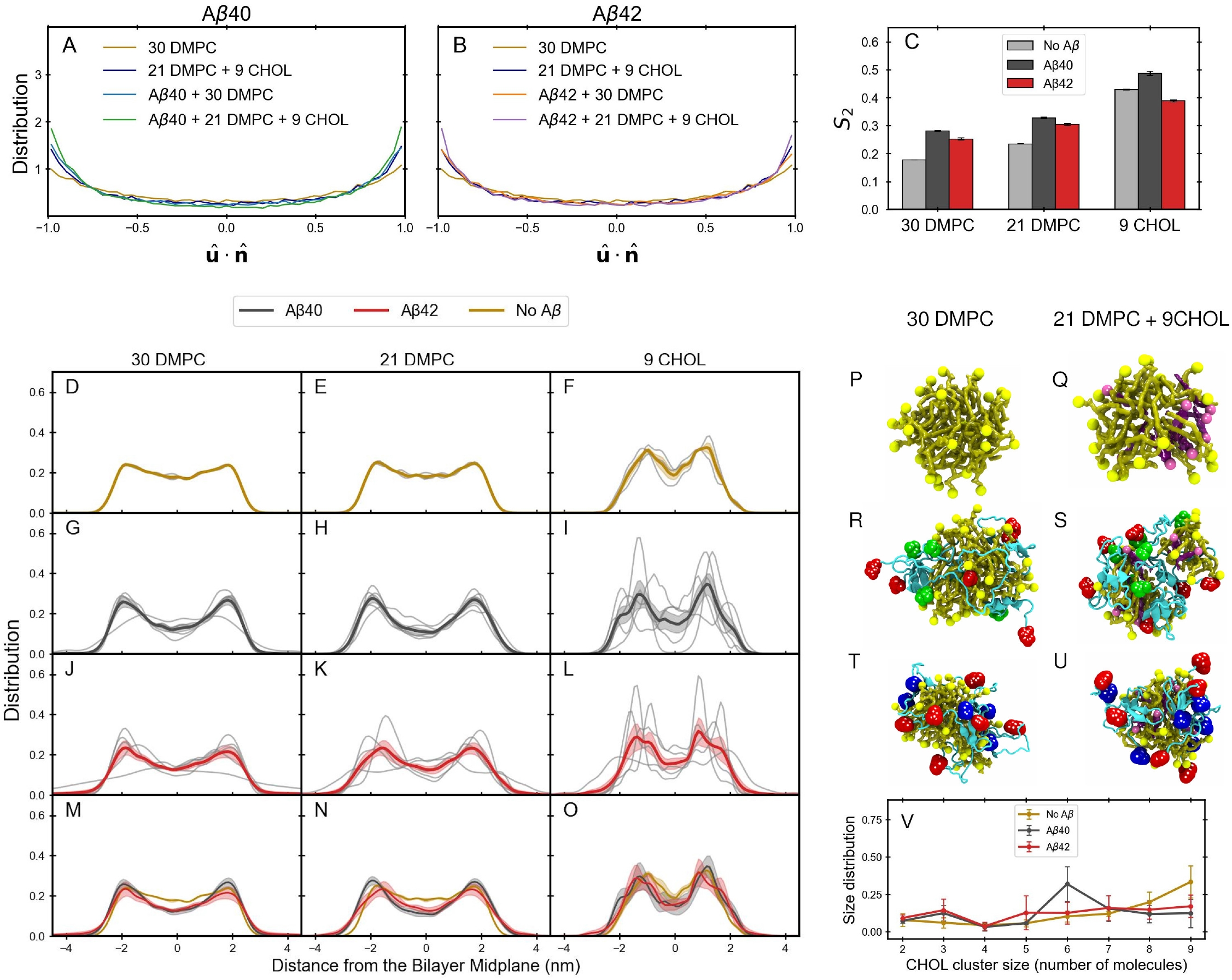
Lipid organization in the absence and presence of A*β*40 and A*β*42. (A,B) Distributions of the projections of lipid molecular directors onto the bilayer normal in the DMPC and DMPC/CHOL systems in the absence and presence of (A) A*β*40 and (B) A*β*42. (C) Average nematic order parameters, *S*_2_, for DMPC and CHOL in the absence of A*β* and in the presence of A*β*40 or A*β*42. (D-O) Lipid headgroup density profiles along the bilayer normal using the DMPC N-headgroup and CHOL O-headgroup. (D-F) Lipids-only systems, (G-I) A*β*40 systems, and (J-L) A*β*42 systems. The three columns correspond to the DMPC system, DMPC in the DMPC/CHOL system, and CHOL in the DMPC/CHOL system, respectively. Thin gray lines represent individual trajectories and thick colored lines represent ensemble averages. (M-O) Corresponding comparisons among the three conditions. All analyses are performed over 900-1000 ns, with error bars and shaded regions representing the standard error of the mean (SEM). (P-U) Representative snapshots of fully assembled systems of (P-Q) lipids only, (R-S) lipids with A*β*40 hexamer, and (T,U) lipids with A*β*42 hexamer for (P,R,T) systems with DMPC lipids and (Q,S,U) systems with DMPC/CHOL mixture created with VMD ^59^. In each snapshot, the normal to the lipid bicelle is vertical. (V) CHOL cluster size distributions in three systems: lipids alone, with A*β*40, and with A*β*42.

Due to a layered structure of a bicelle, a normal to the bicelle can be defined in a meaningful way (see *Computational Methods*). To gain an insight into the morphology of bicelles, we derived for each of the six systems under study a distribution of projections of lipid molecular directors onto the bicelle normal, 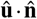. Figure 7A-B shows distinct maxima at 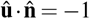and +1 for all six systems, as expected of a system with a preferred orientation. These peaks are less sharp than in the case of the previously reported bilayer-like structure of 100 lipids ^39^, which is consistent with stronger finite size effects in the current case of 30 lipids. We calculated a fraction of lipids whose molecular axes deviate from the normal by no more than 45^*°*^, corresponding to 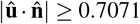. If lipids were arranged into an ideal spherical micelle with radially oriented lipid molecular axes, the corresponding fraction of aligned lipids would be 1− cos 45^*°*^ = 0.293, considering both upper and lower leaflets. Table 1 shows that in the absence of A*β*, fractions of thus aligned lipids are 0.453 and 0.526 for the DMPC and DMPC/CHOL systems, respectively. These results indicate that a significantly larger fraction of lipids are aligned along the bicelle director than in a completely spherical micelle and that CHOL promotes layered structure. This calculation provides a straightforward justification for describing these lipid clusters as bicelles.

Bicelle-like structure is further supported by non-zero DMPC and CHOL nematic order parameters (Figure 7C). Time evolution of DMPC and CHOL nematic order parameters is shown in Figure S7. DMPC lipids that form bicelles in the absence of A*β* are characterized by nematic order parameters *S*_2_ of 0.18 and 0.23 in the DMPC and DMPC/CHOL systems, respectively. Thus, the presence of CHOL increases the nematic order parameter of DMPC lipids. CHOL in the DMPC/CHOL system in the absence of A*β* is characterized by a significantly higher nematic order parameter of 0.43. For comparison, previously reported DMPC and CHOL nematic order parameters of 0.34 and 0.72, respectively, for a DMPC/CHOL/GM1 system of 100 lipids are considerably higher, as expected of a larger, bilayer-like lipid structure ^39^.

Lipid headgroup density profiles with two peaks corresponding to the headgroup regions of the upper and lower leaflet of the layered structure also support the bicelle-like morphology of lipid clusters, although the density between the two peaks remains rather high due to a considerable fraction of lipids aligned away from the normal (Figure 7D-F). In line with the nematic order parameter results, DMPC headgroup profiles exhibit slightly sharper peaks for the DMPC/CHOL system than for the pure DMPC system, whereas the two peaks in the CHOL headgroup profile are the sharpest of the three. The snapshots of bicelles formed in the DMPC and DMPC/CHOL systems are displayed in Figure 7P-Q. It is noteworthy that the three replica MD trajectories of each of these two lipid-only systems produce practically overlapping DMPC headgroup density profiles (Figure 7D-E). In contrast, there is significantly more variability in the CHOL headgroup density profiles (Figure 7F), due to a smaller number of CHOL relative to the number of DMPC molecules.

The average peak-to-peak distance is used to define the thickness of the bicelle, *D*_*B*_, reported in Table 1. In addition, we calculated the area per lipid (APL), which quantifies the average lateral packing of the headgroup regions of a bicelle. Interestingly, the presence of CHOL decreases the DMPC bilayer thickness as well as the area per lipid (APL). The bicelle thickness values for DMPC and CHOL are significantly smaller than those previously reported for the DMPC/CHOL/GM1 system of 100 lipids ^39^. As shown in Table 1, the APL is larger for the system of 30 lipids than for the system of 100 lipids. Both a decrease in the thickness and increase in the APL are consistent with significantly more disordered lipid tails in systems with fewer lipid molecules.

#### 3.2.6 A*β*40 increases the planar organization of a bicelle significantly more than A*β*42

Figure 7A-B show the two peaks at 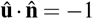 and +1 are sharper when lipids co-assemble with A*β*40 or A*β*42 and are the sharpest for the DMPC/CHOL system, reflecting significantly higher populations of lipids aligned along the normal to a bicelle (Table 1). This effect is considerably stronger for A*β*40 than for A*β*42. In line with these results, the nematic order parameter *S*_2_ for DMPC lipids in both DMPC and DMPC/CHOL systems increases when lipids co-assemble with A*β* (Figure 7C). The A*β*-induced increase in *S*_2_ is significantly higher for A*β*40 than for A*β*42. Interestingly, A*β*40 and A*β*42 co-assembled with DMPC/CHOL system have opposite effects on the nematic order parameter of CHOL, which increases in the presence of A*β*40 and decreases in the presence of A*β*42 relative to the DMPC/CHOL system without A*β*, indicating that unlike A*β*40, A*β*42 disrupts orientational order of CHOL molecules within a bicelle (Figure 7C).

The DMPC and CHOL headgroup density profiles of bicelles formed in the presence of A*β*40 in (Figure 7G-I) or A*β*42 (Figure 7J-L) are sharper than in the absence of A*β* (Figure 7D-F). This result demonstrates that the planar organization of lipid bicelles is enhanced when lipids co-assemble with A*β*. The comparison of DMPC and CHOL headgroup density profiles in Figure 7M-O reveals that these profiles are more pronounced for A*β*40 than for A*β*42, indicating that A*β*40 exerts a stronger ordering effect than A*β*42 on both DMPC and DMPC/CHOL systems. Notably, the trajectory-to-trajectory variability of the DMPC and even more so CHOL headgroup density profile is significant among the five replica trajectories per system, which is a reflection of an intrinsically disordered nature of A*β* even when interacting with lipids. The snapshots of bicelles co-assembled with A*β*40 and A*β*42 in the DMPC and DMPC/CHOL systems are displayed in Figure 7R-U. Notably, CHOL molecules within a bicelle formed in the absence of A*β* tend to cluster together, while A*β* appears to disrupt this CHOL clustering (Figure 7V).

Table 1 shows that the presence of A*β*40 or A*β*42 has no effect on the bicelle thickness in the DMPC system, whereas in the DMPC/CHOL system the DMPC thickness increases in the presence of A*β*40 but not in the presence of A*β*42. This is consistent with the highest planar order associated with the DMPC/CHOL system co-assembled with A*β*40. The presence of A*β*40 and A*β*42 both increases the CHOL thickness within DMPC/CHOL bicelles. A*β* thus significantly increases the APL of a bicelle for both DMPC and DMPC/CHOL systems, which is likely due to a larger size of A*β*-lipid bicelles compared to bicelles formed in the absence of A*β*. Notably, the lipid bilayer-like structures of 100 lipids previously reported for the DMPC/CHOL/GM1 system ^39^ have considerably lower APL values (Table 1), consistent with their more ordered morphology.

#### 3.2.7 A*β*40 interacts with DMPC more avidly than A*β*42

To identify the specific regions of A*β*40 and A*β*42 that may preferentially interact with lipids, we calculated the per-residue contact probabilities of A*β* with DMPC and CHOL (Figure 8), as described in *Computational Methods*. In the DMPC system, A*β*-DMPC contact probability profiles vary considerably along the A*β*40 and A*β*42 sequences (Figure 8A,D). The five highest A*β*40-DMPC contact probabilities are observed for L34, F19, I31, Y10, and F20. Four of these residues belong to the CHC or MHR, indicating an important contribution of these hydrophobic regions to A*β*40-DMPC interactions. The five highest A*β*42-DMPC contact probabilities are observed for F19, L17, M35, V36, and I41, again highlighting the importance of the hydrophobic CHC and MHR, alongside the CTR. The contact probabilities demonstrate that A*β*40 and A*β*42 interact with DMPC lipids in distinct ways. The A*β*40 to A*β*42 contact probability difference in Figure 8G shows that for most residues, these contact probabilities are higher for A*β*40 than for A*β*42. The residues with higher A*β*-DMPC contact probabilities for A*β*42 than for A*β*40 are restricted to a few residues in the CHC, the M35-G37 region, and V39. While hydrophobic regions most strongly participate in A*β*-DMPC interactions, A*β*40 interacts with DMPC lipids more than A*β*42.

**Figure 8.**
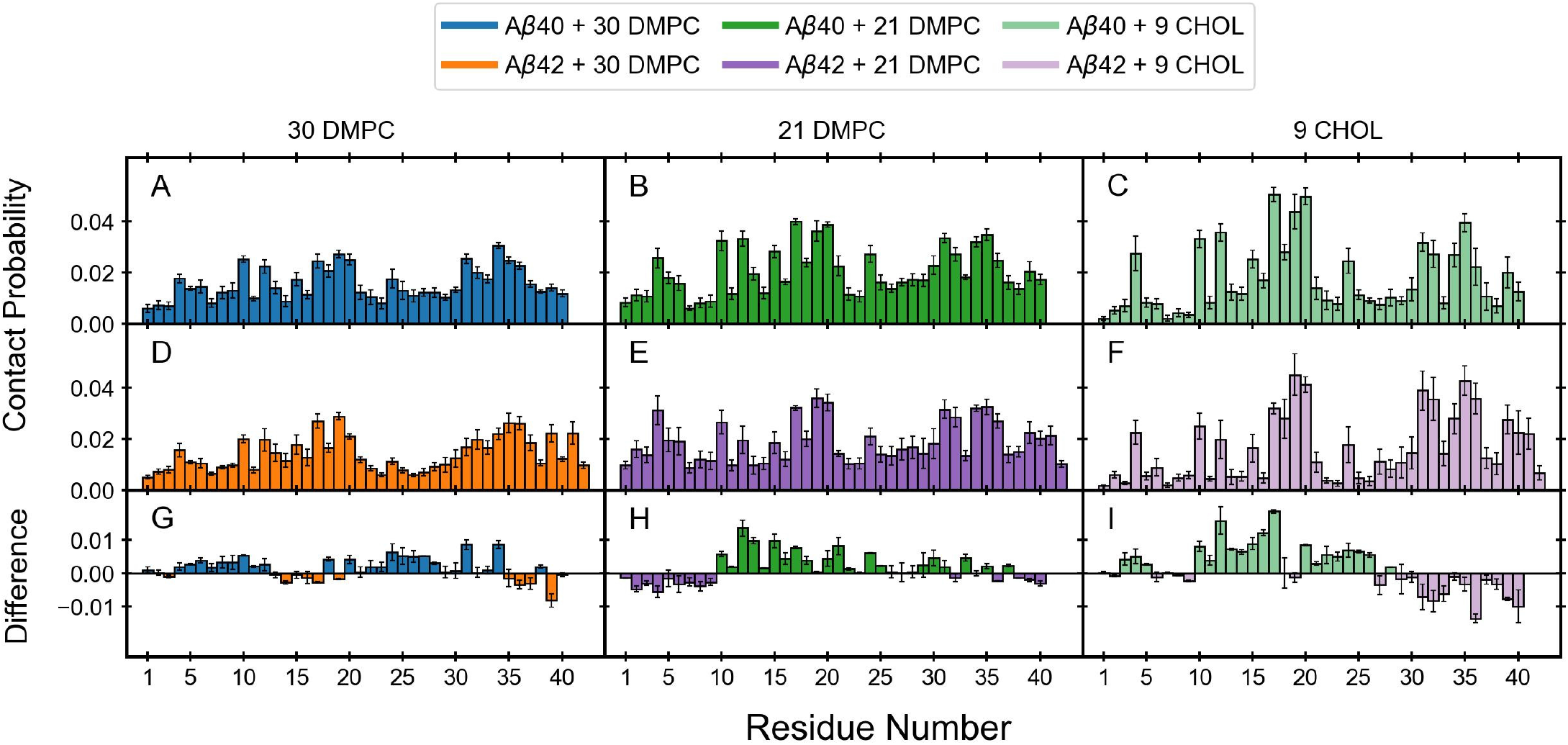
(A,D) Contact probability between DMPC and (A) A*β*40 and (D) A*β*42 in DMPC system. (B,E) Contact probability between DMPC and A*β*, (C,F) CHOL and A*β* in DMPC/CHOL systems. Panels G, H, and I show the A*β*40 - A*β*42 differences for each of the corresponding systems: (G) DMPC, (H-I) DMPC/CHOL. The final values are averaged across 600-1000 ns and over all the trajectories with error bars representing SEM.

#### 3.2.8 CHOL enhances A*β*-DMPC contact probabilities in an isoform-specific manner

Overall, the presence of CHOL in the DMPC/CHOL system increases A*β*-DMPC contact probabilities for both isoforms (Figure 8B,E). This increase is more pronounced for A*β*40 than for A*β*42 (Figure 8B,E). The top five A*β*40-DMPC contact probabilities occur at L17, F20, F19, M35, and I31, whereas the top five A*β*42-DMPC contact probabilities are at F19, F20, M35, L17, and L34. Thus the two hydrophobic regions, CHC and MHR, dominate A*β*-DMPC interactions. Despite this similarity, the A*β*40 to A*β*42 difference probabilities in Figure 8H indicate isoform-dependent character of A*β*-DMPC interactions. A*β*40-DMPC contact probabilities are higher than A*β*42-DMPC contact probabilities for most residues within the broader Y10-M35 region, whereas A*β*42 exhibits higher contact probabilities with DMPC lipids than A*β*40 primarily within the NTR, the adjacent S8-G9 region, at V36 and within the CTR. Thus, the presence of CHOL redistributes A*β*-DMPC interactions, with A*β*40 preferentially interacting with DMPC lipids through a broad central part of the sequence, whereas the stronger A*β*42-DMPC interactions are localized primarily in the vicinity of the two termini.

#### 3.2.9 A*β*-CHOL interactions are dominated by F19 and F20, but also show isoform dependence

In addition to enhancing A*β*-DMPC interactions, CHOL itself interacts strongly with both A*β*40 and A*β*42 through more specific regions or residues. Despite the smaller number of CHOL molecules relative to DMPC lipids in the DMPC/CHOL system, Figure 8C,F indicates that selected residues of both A*β*40 and A*β*42 exhibit A*β*-CHOL contact probabilities that are comparable to or higher than their corresponding A*β*-DMPC contact probabilities. The top five A*β*40-CHOL contact probabilities occur at L17, F20, F19, M35, and V12, whereas the top five A*β*42-CHOL contact probabilities involve F19, M35, F20, I31, and V36. Thus, in both isoforms, F19 and F20 within the CHC exhibit highest affinities for CHOL. However, high V12-CHOL (A*β*40) versus high L17-CHOL (A*β*42) and I31-CHOL (A*β*40) versus M35-CHOL/V36-CHOL (A*β*42) contact probabilities are isoform specific. Figure 8I reveals the A*β*40-CHOL to A*β*42-CHOL difference contact probabilities with a pronounced separation of the regions preferentially interacting with CHOL. Whereas A*β*40-CHOL contact probabilities are higher than A*β*42-CHOL contact probabilities along most of the sequence, up to S26, A*β*42 shows a higher affinity for CHOL than A*β*40 within the N27-A42 region. Thus, CHOL not only alters the A*β*-DMPC contacts but also directly interacts with the two isoforms through distinct regions, whereby A*β*40 interacts with CHOL more than A*β*42 within the N-terminal and central regions while A*β*42 exhibits a stronger preference for CHOL interactions within the C-terminal half of the peptide.

A direct comparison of A*β*-lipid contact probabilities between the two lipid systems in Figure S8A-B supports the above observations. The DMPC contact probabilities, calculated for the DMPC system and subtracted from the corresponding contact probabilities for the DMPC/CHOL system to obtain the A*β*-DMPC contact probability differences, show that CHOL enhances A*β*-DMPC interactions in both A*β*40 and A*β*42 (Figure S8C-D). In contrast, V12, H13, and G33 in A*β*40 and H13, G33, and G37 in A*β*42 show higher A*β*-DMPC contact probabilities in the DMPC system than in the DMPC/CHOL system. Overall, CHOL promotes A*β*-DMPC interactions in both isoforms, with reduced A*β*-DMPC contact probabilities limited to a very small number of residues. The A*β*-lipid contact probabilities, calculated for DMPC lipids in the DMPC system are subtracted from the contact probabilities for CHOL molecules in the DMPC/CHOL system to reveal a more selective pattern of A*β*-CHOL than A*β*-DMPC contacts (Figure S8E-F). A*β*40-CHOL contact probabilities are higher than A*β*40-DMPC contact probabilities for the L17-F20 region within the CHC. A*β*42-CHOL contact probabilities are higher than A*β*42-DMPC contact probabilities within the CHC, particularly for the V18-F20 region, and also for the MHR, including I31, I32, M35, V36, alongside V39 in the CTR.

#### 3.2.10 Unimodal A*β*40-lipid versus bimodal A*β*42-lipid interactions

To examine the energetic modes of A*β*-lipid interactions, Figure 9A-D shows the probability distributions of the A*β*-lipid potential energy per peptide for all four systems with A*β* and lipids. These distributions are Gaussian-like for A*β*40 in both lipid systems, but are non-Gaussian-like for A*β*42, suggesting that while A*β*40 interacts with lipids via a single dominant mode, A*β*42 interactions with lipids may be more complex. In the DMPC system, the mean A*β*40-DMPC potential energy is significantly lower than the mean A*β*42-DMPC potential energy, supporting the conclusion that A*β*40 interacts with DMPC lipids more avidly than A*β*42. Interestingly, in the presence of CHOL in the DMPC/CHOL system, the mean A*β*40-lipid potential energy significantly increases relative to that in the DMPC system, while the corresponding increase in the mean A*β*42-lipid potential energy is not statistically significant. While both A*β*40-lipid and A*β*42-lipid potential energy distributions in the DMPC system exhibit a single peak, the A*β*42-lipid potential energy is asymmetric, suggesting that even in pure DMPC lipids, there may be two A*β*42-lipid interaction modes. This conjecture is further supported by an observation that in the DMPC/CHOL system, A*β*42-lipid potential energy distribution is clearly bimodal. Thus, A*β*42 in the DMPC/CHOL system interacts with lipids via two distinct modes.

**Figure 9.**
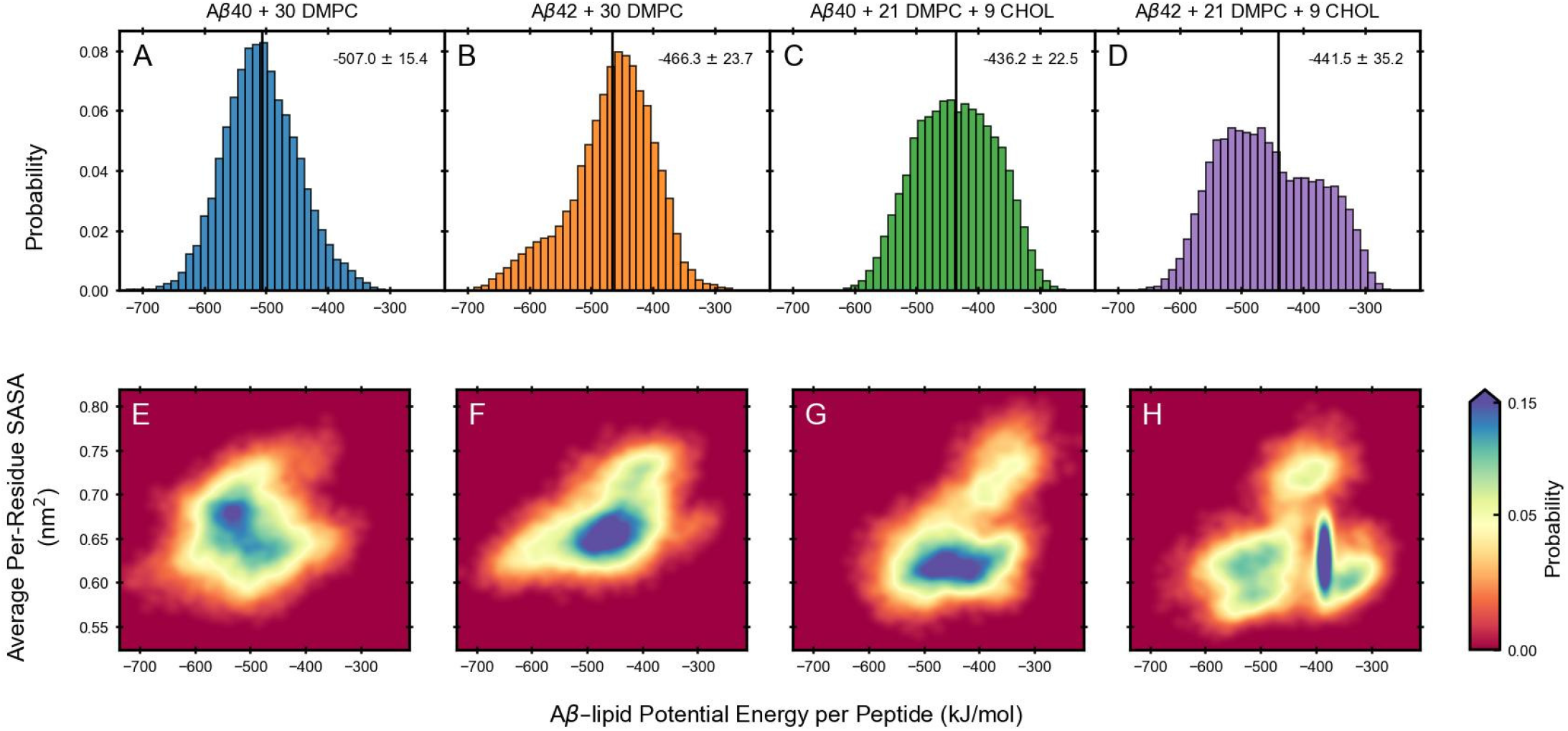
A*β*-lipid potential energy and total Solvent-Accessible Surface Area (SASA) of A*β*40 and A*β*42 in the DMPC and DMPC/CHOL systems. (A-D) Probability distributions of the A*β*-lipid potential energy per peptide in the DMPC (A,B) and DMPC/CHOL (C,D) systems with A*β*40 (A,C) or A*β*42 (B,D). Vertical black lines indicate the mean potential energies, with corresponding mean *±*SEM values shown in each panel. (E-H) Two dimensional probability distributions of the A*β*-lipid potential energy per peptide and total SASA per residue for the corresponding systems in A-D. All analyses were performed over 900-1000 ns of the five replica trajectories per system.

#### 3.2.11 Competition between A*β*-A*β* and A*β*-lipid interactions

We here asked whether the extent of A*β* solvent exposure can help identify the origin of distinct A*β*-lipid interactions of the two isoforms. To this end, we derive two-dimensional (2D) probability distributions using the A*β*-lipid potential energy per peptide and average per-residue SASA as the two coordinates. 2D distributions in Figure 9E-F demonstrate that in the DMPC system, low A*β*40-DMPC potential energies are associated with a rather high average per-residue SASA whereas average per-residue SASA of A*β*42 is higher and associated with higher A*β*42-DMPC potential energies than in the case of A*β*40. 2D distributions in Figure 9G-H, on the other hand, indicate that in the DMPC/CHOL system, relative to the DMPC system, an average per-residue SASA is significantly decreased for both isoforms. Thus, CHOL induces a more efficient shielding from the solvent in both isoforms. Importantly, there is a clear difference between A*β*40 and A*β*42 probability distributions in the DMPC/CHOL system, i.e. A*β*42 but not A*β*40 is associated with two distinct basins that are separated along the A*β*-lipid potential energy coordinate. Interestingly, the dominant basin corresponds to high A*β*42-lipid potential energies, suggesting that the dominant A*β*42-lipid interaction mode is associated with a narrowly defined range of relatively high A*β*-lipid interaction energies.

To examine A*β*40 versus A*β*42 difference in A*β*-lipid interaction modes in more detail, we derived 2D probability distributions with the A*β*-lipid potential energy per peptide and average per-residue SASA as the two coordinates for each replica trajectory of each of the four systems (Figure S9). While all systems clearly display some trajectory-to-trajectory variability, A*β*42 co-assembly with lipids in the DMPC/CHOL system exhibits the most pronounced trajectory-dependent behavior, whereby trajectories 1 and 4 predominantly sample the narrow high potential energy region and therefore contribute to the high potential energy peak in the bimodal distribution in Figure 9D. Because the trajectory-specific populations do not show any separation along the average per-residue SASA coordinate, we then ask if these two interaction modes might reflect a competition between A*β*42-A*β*42 and A*β*-lipid interactions. We posit that the total A*β* potential energy, i.e. the sum of A*β*-A*β* and A*β*-lipid potential energies, is partitioned among the two contributions in isoform- and lipid system-specific ways. To test this conjecture, we calculate average A*β*-A*β* potential energies for a comparison with average A*β*-lipid potential energies for each of the five replica trajectories of each of the four systems. The results in Table 2 indeed demonstrate that there is a competition between A*β*-A*β* and A*β*-lipid interactions in all four systems. For A*β*42 in the DMPC/CHOL system, trajectories 1 and 4, which contribute to the higher A*β*-lipid energy population, are characterized by the highest A*β*42-A*β*42 percentage contribution and the lowest A*β*42-lipid percentage contribution to the total potential energy, indicating that the high A*β*42-lipid potential energy peak in Figure 9D is due to A*β*42-A*β*42 self-assembly which dominates over the A*β*42-lipid co-assembly. While this competition exists in the other three systems as well, it leads to two different A*β*-lipid interaction modes only for A*β*42 in the DMPC/CHOL system.

**Table 2.**
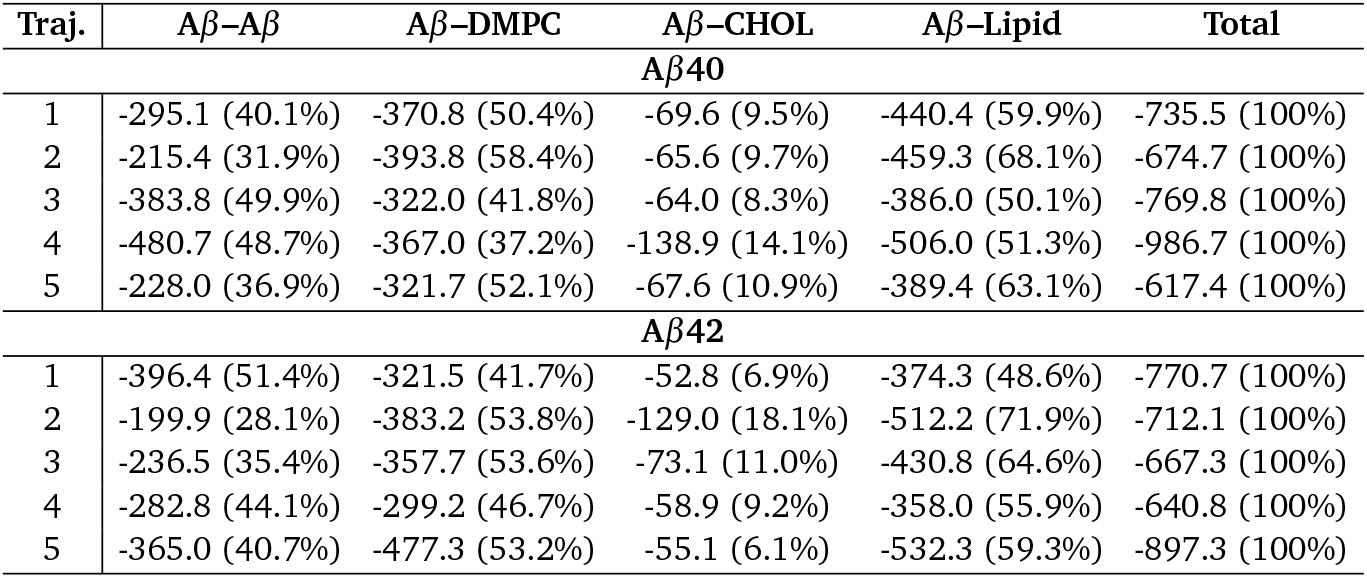
Interaction energies for A*β*40 and A*β*42 across five independent trajectories. Values in parentheses represent the percentage contribution relative to the total energy of the corresponding trajectory.

| Traj. | Aβ-Aβ | Aβ-DMPC | Aβ-CHOL | Aβ-Lipid | Total |
| --- | --- | --- | --- | --- | --- |
| **Aβ40** |  |  |  |  |  |
| 1 | -295.1 (40.1%) | -370.8 (50.4%) | -69.6 (9.5%) | -440.4 (59.9%) | -735.5 (100%) |
| 2 | -215.4 (31.9%) | -393.8 (58.4%) | -65.6 (9.7%) | -459.3 (68.1%) | -674.7 (100%) |
| 3 | -383.8 (49.9%) | -322.0 (41.8%) | -64.0 (8.3%) | -386.0 (50.1%) | -769.8 (100%) |
| 4 | -480.7 (48.7%) | -367.0 (37.2%) | -138.9 (14.1%) | -506.0 (51.3%) | -986.7 (100%) |
| 5 | -228.0 (36.9%) | -321.7 (52.1%) | -67.6 (10.9%) | -389.4 (63.1%) | -617.4 (100%) |
| **Aβ42** |  |  |  |  |  |
| 1 | -396.4 (51.4%) | -321.5 (41.7%) | -52.8 (6.9%) | -374.3 (48.6%) | -770.7 (100%) |
| 2 | -199.9 (28.1%) | -383.2 (53.8%) | -129.0 (18.1%) | -512.2 (71.9%) | -712.1 (100%) |
| 3 | -236.5 (35.4%) | -357.7 (53.6%) | -73.1 (11.0%) | -430.8 (64.6%) | -667.3 (100%) |
| 4 | -282.8 (44.1%) | -299.2 (46.7%) | -58.9 (9.2%) | -358.0 (55.9%) | -640.8 (100%) |
| 5 | -365.0 (40.7%) | -477.3 (53.2%) | -55.1 (6.1%) | -532.3 (59.3%) | -897.3 (100%) |

## 4 Discussion and Conclusions

This experiment-driven MD study focuses on A*β*40 and A*β*42 co-assembly with DMPC lipids and the effect of CHOL on *Aβ*-DMPC co-assembly pathways and interaction modes. We examine kinetics of A*β*40 and A*β*42 fibril formation at different PL ratios using ThT fluorescence spectroscopy to identify the PL ratio of 1:5, which affects the fibril formation rate the most. Experimental results reveal that at this PL ratio, DMPC lipids significantly increase the lag time for A*β*40 but not A*β*42 fibril formation, whereas the presence of CHOL in the DMPC/CHOL system further slows down A*β*40 fibril formation and, to a lesser extent, also A*β*42 fibril formation.

Positing that these contrasting effects of lipids on A*β*40 and A*β*42 fibril formation stem from distinct A*β*-lipid interaction modes of the two isoforms at early co-assembly stages, we perform extensive MD simulations to elucidate atomistic details of early stages of co-assembly of six A*β*40/A*β*42 peptides with 30 DMPC lipids (the DMPC system) and 21 DMPC and 9 CHOL molecules (the DMPC/CHOL system) using the PL ratio of 1:5, to match the experimental systems. We acquire 5 replica MD trajectories, each 1*µ*s long, for each A*β*-containing system to account for the IDP nature of A*β* peptides that requires enhanced sampling of the configurational space ^19^. A*β*-containing systems converge to a quasi steady-state by 600 ns and the systems are dominated by A*β* hexamers. Most A*β*-centered analysis is performed using frames within 600-1000 ns of each relevant trajectory. While lipid-only control systems assemble into a single lipid cluster within the first 200 ns of simulation time, in the presence of A*β*, this process takes significantly longer. Thus, the lipid-centered analysis uses 900-1000 ns of each relevant trajectory.

Our MD results demonstrate that while in the lipid-free systems, A*β*42 hexamer propensity is higher than A*β*40 hexamer propensity, in the presence of lipids the opposite is true. The presence of CHOL in the DMPC/CHOL system results in increased stability of hexamers of both isoforms, indicating that the oligomer size distribution is both isoform- and lipid system-specific. Lipids also affect the average A*β*40 secondary structure by increasing the average *β*-strand, turn, and helical content at the expense of statistical coil content. The DMPC/CHOL system exerts a stronger effect on the A*β*40 secondary structure than the DMPC system. In contrast, the average A*β*42 secondary structure content is almost unaffected by the lipids. Interestingly, A*β*42 is characterized by higher average *β*-strand content than A*β*40 in the absence and presence of lipids, in line with the ability of the longer isoform to aggregate into fibrils faster than the shorter isoform. Lipids also modify the tertiary and quaternary structure of A*β* hexamers in an isoform- and lipid system-specific manner.

All lipid-containing systems assemble into a planar micelles (bicelle), which is distinguishable from an ideal spherical micelle by hosting a significantly larger fraction of lipid molecules aligned toward the normal to the two leaflets of the bicelle (at least 0.45) than expected for a micelle (0.29). The planar aspect of bicelles made of 30 lipids observed here is less obvious than in the case of 100 lipids reported by Pandey and collaborators ^39^, yet it can nonetheless be quantified in a meaningful way by DMPC and CHOL nematic order parameters, headgroup density profiles, thickness along the normal of the bicelle, and APL. Our lipids-only bicellar structures are characterized by DMPC nematic order parameters of 0.18 and 0.23 in the A*β*-free DMPC and DMPC/CHOL systems, respectively, and the CHOL order parameter of 0.43 in the DMPC/CHOL system. The measured DMPC and CHOL thickness of bicelles made of 30 lipids is smaller and the APL larger than the thickness and APL of previously reported bilayer-like structures formed by 100 lipids ^39^, which is consistent with more disordered lipid tails in smaller systems. In line with the previous MD study with 100 lipids (68 DMPC, 30 CHOL, and 2 GM1), we note that CHOL has a tendency to cluster within bicelles, although not all CHOL molecules form a single cluster like in the case of 100 lipids, likely due to a smaller number of lipids.

Perhaps the most intriguing observation is that co-assembly of both A*β*40 and A*β*42 with lipids results in increased planar organization of DMPC lipids as reflected in the increased population of lipids aligned along the normal to the bicelle, increased DMPC nematic order, and sharper headgroup lipid density profiles. Notably, A*β*40 exerts a much stronger lipid ordering effect than A*β*42. In the DMPC/CHOL system, A*β*40 increases the CHOL nematic order, while A*β*42 exerts the opposite effect by considerably decreasing the nematic order of CHOL. The analysis of A*β*-lipid contact probabilities shows that A*β*40 interacts with lipids more avidly than A*β*42 in both lipid systems. In the DMPC/CHOL system, the presence of CHOL results in increased A*β*-DMPC contact probabilities for both isoforms. While A*β*-CHOL interactions are dominated by F19 and F20 in both isoforms, A*β*40 tends to interact with CHOL more through the N-terminal half, while A*β*42 interacts with CHOL more through the C-terminal half of the peptide sequence. The analysis of A*β*-lipid and A*β*-A*β* potential energy distributions indicates that in the DMPC/CHOL system, A*β*42 displays two interaction modes with lipids associated with a bimodal A*β*42-lipid potential energy distribution, whereby the peaks at the lower and higher potential energy correspond to A*β*42-lipid association and A*β*42-A*β*42 self assembly, respectively. While a competition between A*β*-A*β* and A*β*-lipid interactions must be present in all four systems, it leads to clearly separated modes of A*β*-lipid interaction only for A*β*42 in the DMPC/CHOL system.

The above findings lead to the following understanding of our experimental results that elucidate the stronger effect of lipids on A*β*40 than A*β*42 and the peculiar role of CHOL that modulates A*β*-lipid interactions. The analysis of MD simulations indicates that A*β*40-lipid interactions are stronger than A*β*42-lipid interactions, which gives a plausible explanation of why lipids perturb A*β*40-A*β*40 interactions, leading to an increased lag time for fibril formation, but have a lesser effect on A*β*42-A*β*42 interactions. Moreover, CHOL facilitates stronger A*β*-DMPC interactions in the DMPC/CHOL system for both isoforms, providing further insights into why the presence of CHOL in our experiments further delays A*β*40 fibril formation and also increases the lag time of A*β*42 fibril formation. Thus, a competition between A*β*-lipid association and A*β*-A*β* self-assembly drives the isoform-specific effects of lipids on A*β* fibril formation. Our MD results can also be used to elucidate the emergence of two A*β*42-lipid interaction modes in the DMPC/CHOL system. The critical role of the C-terminal region in A*β*42-A*β*42 oligomer formation is well known. Our MD results indicate that this same region is also more strongly involved in A*β*42-CHOL interactions, which implies that A*β*42-A*β*42 hexamer conformational ensemble is incompatible with A*β*42-CHOL conformational ensemble, leading to the two distinct, mutually-exclusive interaction modes.

Finally, our MD findings also reveal that although A*β*40-lipid interactions are stronger than A*β*42-lipid interactions, the resulting lipid bicellar structure is more ordered in the presence of A*β*40 hexamers than in the presence of A*β*42 hexamers. The contrasting isoform-specific effects on CHOL are particularly intriguing. A previous study by Pandey and collaborators revealed a strong preference of A*β*42 dimers but less so of A*β*42 monomers for CHOL ^39^. These combined MD findings are thus in agreement with the most recent experimental study by Baral and collaborators, who reported an intriguing ability of A*β*42 assemblies to bind to and remove CHOL from a membrane, changing its mechanical properties and potentially inducing membrane damage ^62^. Our MD findings show that A*β*40 hexamers are significantly less capable of interacting with CHOL than A*β*42 assemblies, which should result in their decreased capacity to remove CHOL from a membrane when compared to A*β*42 assemblies, a prediction that is amenable to experimental testing.

## Author contributions

M.L. and B.U. designed the study, T.O. and M.L. collected ThT data, R.P. performed and analyzed the simulations, and R.P., M.L., and B.U. prepared the manuscript.

## Data availability

The data produced in this study are available on request from the corresponding author.

## Acknowledgements

This work has been supported by the National Institutes of Health grant NS139178 to M.L. MD simulations were performed on hardware supported by Drexel’s University Research Computing Facility and on Bridges-2 at Pittsburgh Supercomputing Center through allocation BIO250006 from the Advanced Cyberinfrastructure Coordination Ecosystem: Services & Support (ACCESS) program, which is supported by National Science Foundation grants #2138259, #2138286, #2138307, #2137603, and #2138296 ^63^.

## References

[1] D. J. Selkoe, Physiological Reviews, 2001, 81, 741–766.

[2] J. A. Hardy and G. A. Higgins, Science, 1992, 256, 184–185.

[3] J. Hardy and D. J. Selkoe, Science, 2002, 297, 353–356.

[4] K. N. Dahlgren, A. M. Manelli, W. B. Stine, L. K. Baker, G. A. Krafft and M. J. LaDu, J. Biol. Chem., 2002, 277, 32046–32053.

[5] G. Bitan, M. D. Kirkitadze, A. Lomakin, S. S. Vollers, G. B. Benedek and D. B. Teplow, Proc. Natl. Acad. Sci. U. S. A., 2003, 100, 330–335.

[6] L. M. Shaw, H. Vanderstichele, M. Knapik-Czajka, C. M. Clark, P. S. Aisen, R. C. Petersen, K. Blennow, H. Soares, A. Simon, P. Lewczuk, R. Dean, E. Siemers, W. Potter, V. M. Y. Lee, J. Q. Trojanowski and A. D. N. Initi, Ann. Neurol., 2009, 65, 403–413.

[7] H. B. Pollard, E. Rojas and N. Arispe, in Alzheimer’s disease: Amyloid precusor proteins, signal transduction, and neuronal transplantation, ed. R. M. Nitsch, J. H. Growdon, S. Corkin and R. J. Wurtman, 1993, vol. 695 of Ann. N. Y. Acad. Sci., pp. 165–168.

[8] A. Quist, I. Doudevski, H. Lin, R. Azimova, D. Ng, B. Frangione, B. Kagan, J. Ghiso and R. Lal, Proc. Natl. Acad. Sci. U. S. A., 2005, 102, 10427–10432.

[9] G. M. Shankar, B. L. Bloodgood, M. Townsend, D. M. Walsh, D. J. Selkoe and B. L. Sabatini, J. Neurosci., 2007, 27, 2866–2875.

[10] A. Demuro, I. Parker and G. E. Stutzmann, J. Biol. Chem., 2010, 285, 12463–12468.

[11] M. Arbel-Ornath, E. Hudry, J. R. Boivin, T. Hashimoto, S. Takeda, K. V. Kuchibhotla, S. Hou, C. R. Lattarulo, A. M. Belcher, N. Shakerdge, P. B. Trujillo, A. Muzikansky, R. A. Betensky, B. T. Hyman and B. J. Bacskai, Mol. Neurodegener., 2017, 12, year.

[12] T. Yasumoto, Y. Takamura, M. Tsuji, T. Watanabe-Nakayama, K. Imamura, H. Inoue, S. Nakamura, T. Inoue, A. Kimura, S. Yano, H. Nishijo, Y. Kiuchi, D. B. Teplow and K. Ono, FASEB J., 2019, 33, 9220–9234.

[13] M. Kawahara, M. Kato-Negishi and K.-I. Tanaka, Molecules, 2020, 25, 1441.

[14] G. Fani, B. Mannini, G. Vecchi, R. Cascella, C. Cecchi, C. M. Dobson, M. Vendruscolo and F. Chiti, ACS Chem. Neurosci., 2021, 12, 766–781.

[15] T. L. Williams and L. C. Serpell, FEBS J., 2011, 278, 3905–3917.

[16] H. M. Brothers, M. L. Gosztyla and S. R. Robinson, Front. Aging Neurosci., 2018, 10, 118.

[17] M. R. White, R. Kandel, I.-N. Hsieh, X. De Luna and K. L. Hartshorn, PLoS One, 2018, 13, e0194001.

[18] D. K. Kumar, S. H. Choi, K. J. Washicosky, W. A. Eimer, S. Tucker, J. Ghofrani, A. Lefkowitz, G. McColl, L. E. Goldstein, R. E. Tanzi and R. D. Moir, Sci. Transl. Med., 2016, 8, 340ra72.

[19] J.-E. Shea and B. Urbanc, Curr. Topics Med. Chem., Special Issue: Protein Misfolding in Conformational Disorders, Guest Ed. Cláudio M. Gomes, 2012, 12, 2596–2610.

[20] B. Urbanc, Israel J. Chem., 2017, 57, 651–664.

[21] B. Urbanc, J. Phys. Chem. B, 2021, 125, 1307–1316.

[22] B. Urbanc, L. Cruz, S. Yun, S. V. Buldyrev, G. Bitan, D. B. Teplow and H. E. Stanley, Proc. Natl. Acad. Sci. U. S. A., 2004, 101, 17345–17350.

[23] N. G. Sgourakis, Y. Yan, S. A. McCallum, C. Wang and A. E. Garcia, J. Mol. Biol., 2007, 368, 1448–1457.

[24] B. Urbanc, M. Betnel, L. Cruz, G. Bitan and D. B. Teplow, J. Am. Chem. Soc., 2010, 132, 4266–4280.

[25] B. Barz and B. Urbanc, PLoS One, 2012, 7, e34345.

[26] D. J. Rosenman, C. Wang and A. E. Garcia, J. Phys. Chem. B, 2016, 120, 259–277.

[27] M. J. Voelker, B. Barz and B. Urbanc, J. Chem. Theory Comput., 2017, 13, 4567–4583.

[28] R. Pandey and B. Urbanc, Biomolecules, 2024, 14, 774.

[29] C. H. Davis and M. L. Berkowitz, Biophys. J., 2009, 96, 785–797.

[30] C. H. Davis and M. L. Berkowitz, J. Phys. Chem. B, 2009, 113, 14480–14486.

[31] A. M. Brown and D. R. Bevan, Biophys. J., 2016, 111, 937–949.

[32] W. Feng, H. Lei, J. Si and T. Zhang, PLoS One, 2017, 12, year.

[33] H. Fatafta, M. Khaled, M. C. Owen, A. Sayyed-Ahmad and B. Strodel, Proc. Natl. Acad. Sci. U. S. A., 2021, 118, e2106210118.

[34] B. Wang and C. Guo, ACS Chem. Neurosci., 2022, 13, 2709–2718.

[35] O. Press-Sandler and Y. Miller, Protein Sci., 2022, 31, year.

[36] H. Fatafta, B. Kav, B. F. Bundschuh, J. Loschwitz and B. Strodel, Biophys. Chem., 2022, 280, 106700.

[37] A. Chakravorty, S. D. McCalpin, B. R. Sahoo, A. Ramamoorthy and C. L. Brooks III, J. Phys. Chem. Lett., 2022, 13, 9303–9308.

[38] B. Andrews, T. Ruggiero and B. Urbanc, Phys. Chem. Chem. Phys., 2023, 25, 2566–2583.

[39] R. Pandey, T. Ruggiero, B. Andrews and B. Urbanc, ACS. Chem. Neurosci., 2026, 17, 2469–2485.

[40] H. J. C. Berendsen, D. van der Spoel and R. van Drunen, Comput. Phys. Commun., 1995, 91, 43–56.

[41] E. Lindahl, B. Hess and D. van der Spoel, J. Mol. Model., 2001, 7, 306–317.

[42] D. V. D. Spoel, E. Lindahl, B. Hess, G. Groenhof, A. E. Mark and H. J. C. Berendsen, J. Comput. Chem., 2005, 26, 1701–1718.

[43] B. Hess, C. Kutzner, D. van der Spoel and E. Lindahl, J. Chem. Theory Comput., 2008, 4, 435–447.

[44] S. Pronk, S. Pall, R. Schulz, P. Larsson, P. Bjelkmar, R. Apostolov, M. R. Shirts, J. C. Smith, P. M. Kasson, D. van der Spoel, B. Hess and E. Lindahl, Bioinformatics, 2013, 29, 845–854.

[45] S. Pall, M. J. Abraham, C. Kutzner, B. Hess and E. Lindahl, Solving software challenges for exascale, 2015, pp. 3–27.

[46] M. J. Abraham, T. Murtola, R. Schulz, S. Páll, J. C. Smith, B. Hess and E. Lindahl, SoftwareX, 2015, 1-2, 19 – 25.

[47] A. D. MacKerell, D. Bashford, M. Bellott, R. L. Dunbrack, J. D. Evanseck, M. J. Field, S. Fischer, J. Gao, H. Guo, S. Ha, D. Joseph-McCarthy, L. Kuchnir, K. Kuczera, F. T. K. Lau, C. Mattos, S. Michnick, T. Ngo, D. T. Nguyen, B. Prodhom, W. E. Reiher, B. Roux, M. Schlenkrich, J. C. Smith, R. Stote, J. Straub, M. Watanabe, J. Wiorkiewicz-Kuczera, D. Yin and M. Karplus, J. Phys. Chem. B, 1998, 102, 3586–3616.

[48] MacKerell, A. D. and Feig, M. and Brooks, C. L., J. Comput. Chem., 2004, 25, 1400–1415.

[49] R. B. Best, X. Zhu, J. Shim, P. E. M. Lopes, J. Mittal, M. Feig and A. D. MacKerell, Jr., J. Chem. Theory Comput., 2012, 8, 3257–3273.

[50] J. Huang, S. Rauscher, G. Nawrocki, T. Ran, M. Feig, B. L. de Groot, H. Grubmüller and A. D. MacKerell, Jr., Nat. Methods, 2017, 14, 71–73.

[51] J. B. Klauda, R. M. Venable, J. A. Freites, J. W. O’Connor, D. J. Tobias, C. Mondragon-Ramirez, I. Vorobyov, A. D. MacKerell, Jr. and R. W. Pastor, J. Phys. Chem. B, 2010, 114, 7830–7843.

[52] R. W. Pastor and A. D. MacKerell, Jr., J. Phys. Chem. Lett., 2011, 2, 1526–1532.

[53] R. M. Venable, A. J. Sodt, B. Rogaski, H. Rui, E. Hatcher, A. D. MacKerell, Jr., R. W. Pastor and J. B. Klauda, Biophys. J., 2014, 107, 134–145.

[54] S. Pall and B. Hess, Comput. Phys. Commun., 2013, 184, 2641–2650.

[55] S. Nosé, J. Chem. Phys., 1984, 81, 511–519.

[56] W. G. Hoover, Phys. Rev. A, 1985, 31, 1695–1697.

[57] M. Parrinello and A. Rahman, J. Appl. Phys., 1981, 52, 7182–7190.

[58] M. Heinig and D. Frishman, Nucleic Acids Res., 2004, 32, W500–W502.

[59] W. Humphrey, A. Dalke and K. Schulten, J. Mol. Graphics, 1996, 14, 33–38.

[60] R. T. McGibbon, K. A. Beauchamp, M. P. Harrigan, C. Klein, J. M. Swails, C. X. Hernandez, C. R. Schwantes, L.-P. Wang, T. J. Lane and V. S. Pande, Biophys. J., 2015, 109, 1528–1532.

[61] P. Virtanen, R. Gommers, T. E. Oliphant, M. Haberland, T. Reddy, D. Cournapeau, E. Burovski, P. Peterson, W. Weckesser, J. Bright et al., Nature Methods, 2020, 17, 261–272.

[62] R. Baral, R. van Deventer and Y. L. Lyubchenko, J. Neurochem., 2026, 170, e70380.

[63] T. J. Boerner, S. Deems, T. R. Furlani, S. L. Knuth and J. Towns, in Practice and experience in advanced research computing: Computing for the common good, 2023, pp. 173–176.

