## Supporting information for "Distinct A*β*40 and A*β*42 co-assembly pathways with free lipids: Experiment-driven MD study^†^"

### Supplementary Information: Distinct A$\beta$40 and A$\beta$42 co-assembly pathways with free lipids: Experiment-driven MD study

Rachit Pandey[a], Tarek Olabi[b], Myungwoon Lee[c], Brigita Urbanc[a*]

[a]Physics Department, Drexel University, 3141 Chestnut St, Philadelphia, 19104, PA, USA.
[b]Department of Biology, Drexel University, 3141 Chestnut St, Philadelphia, 19104, PA, USA.
[c]Department of Chemistry, Drexel University, 3141 Chestnut St, Philadelphia, 19104, PA, USA.

#### Supplementary Molecular Dynamics Results

##### Effect of lipids on the tertiary structure of A$\beta$ hexamers

To investigate how lipids affect tertiary structure of A$\beta$ hexamers, we calculated the intramolecular contact maps for both A$\beta$40 and A$\beta$42 in the absence and presence of DMPC and DMPC/CHOL lipids as described in *Methods*.

In the absence of lipids, the most prominent long-range intramolecular contacts in A$\beta$40 form between regions H6-E11 and K28-I32 (Figure 5A). Additional tertiary contacts form between regions K16-A21 and V36-G38 as well as between regions F20-V24 and G29-A30 (Figure 5A). The top five most probable contacts are: G9-A30, E11-I32, G9-G29, Y10-I31, and G9-I31. The dominance of contacts connecting regions G9-E11 with G29-I32 indicates that the most prominent albeit transient tertiary A$\beta$40 structure arises from long-range contacts between the N-terminal region G9-E11 and MHR. A$\beta$42 in the absence of lipids forms the most prominent contacts between regions G9-H14 and G33-L34, between regions L17-F20 and A30-M35, and between regions A21-V24 and N27-A30 (Figure 5B). The five strongest A$\beta$42 tertiary contacts are: L17-I32, F19-A30, H14-L34, F20-G29, and H14-G33. A$\beta$42 thus adopts a transient tertiary conformation, in which the CHC and neighboring residues are proximate to the MHR, thereby forming a loop-like structure within region E22-A30. While this loop-like structure is present also in A$\beta$40, it is significantly less prominent than in A$\beta$42. Consistent with these observation, the A$\beta$42 to A$\beta$40 difference map shows that contacts L17-I32, F19-A30, H14-L34, F20-G29, and H14-G33 are all more probable in A$\beta$42 than in A$\beta$40 (Figure 5C). In comparison, contacts E11-L34, G9-I32, D23-A30, Y10-G33, and G9-I31 in A$\beta$40 are more probable than in A$\beta$42. Thus, in the absence of lipids, transient tertiary structure A$\beta$40 is dominated by contacts between region G9-E11 and the MHR, whereas A$\beta$42 favors a more centrally organized tertiary contact network connecting the CHC with the MHR, resulting in a more prominent loop-like feature in region E22-A30.

DMPC lipids substantially reorganize tertiary structure of A$\beta$40 (Figure 5D). A broader tertiary contact network connects regions F20-V24 and K28-A30. Additional prominent tertiary contacts form between regions V18-A21 and I32-G33, between regions H14-V18 and V36-G38, and between regions A2-R5 and V18-A21. The five strongest A$\beta$40 tertiary contacts are: F20-G29, E11-K16, F20-I31, L17-E22, and F20-A30. The predominance of contacts involving F20 indicates that the presence of DMPC lipids shifts the tertiary organization of A$\beta$40 from contacts between the region G9-E11 and MHR (which are prominent in the absence of lipids) toward an F20-centered network connecting the CHC with region G29-I31. The most probable A$\beta$42 tertiary contacts in the DMPC system occur between regions E11-K16 and V12-L17 and between regions F19-V24 and N27-A30 (Figure 5E). Additional long-range contacts connect the CHC with region G38-I41 and the MHR with the CTR. The five strongest contacts are: L34-V40, E11-K16, V12-L17, F20-G29, and F19-V39. These contacts indicate that the presence of DMPC lipids promotes an A$\beta$42 tertiary organization involving both contacts between region E11-L17 and the central portion of the sequence and contacts between the CHC and the MHR as well as between the MHR and the CTR. The A$\beta$42 to A$\beta$40 difference map further elucidates the ways in which the presence of DMPC affects A$\beta$ tertiary structures in an isoform-dependent manner (Figure 5F). The five largest A$\beta$42-favored difference contacts are: L34-V40, E11-K16, V12-L17, F20-G29, and E22-N27, followed by G33-I41 and G33-A42. The five top A$\beta$40-favored difference contacts are R5-F20, E11-F20, F20-I31, F20-G33, and G9-E22. Thus, in the presence of DMPC, A$\beta$40 favors F20-centered tertiary contacts, whereas A$\beta$42 favors contacts involving region E11-L17 together with contacts between the MHR and CTR.

In the DMPC/CHOL system, A$\beta$40 exhibits prominent long-range contacts between regions H6-Y10 and G25-G29 and between regions G9-E11 and A30-M35 (Figure 5G). Additional contacts form between regions F20-V24 and G25-G29 and between regions K16-A21 and G37-G38. The top five contacts with the highest probability to form are: G9-A30, E11-V18, G9-F20, G9-G29, and E11-I32. The predominance of contacts involving region G9-E11 indicates that the tertiary organization of A$\beta$40 in the DMPC/CHOL system is largely supported by interactions connecting this sequence segment with the CHC and region G29-I32. A$\beta$42 exhibits prominent long-range contacts between regions Q15-F20 and G29-G33 and between regions V12-H14 and G33-M35 (Figure 5H). Additional contacts form between regions H6-G9 and K16-L17 and between regions L17-F20 and I31-G33. The top five most probable contacts are: R5-V18, D7-K16, L17-I32, F19-A30, and H14-L34, revealing that in the DMPC/CHOL system, A$\beta$42 tertiary organization involves interactions of the NTR with residues K16-V18 within the CHC, alongside the contacts connecting the CHC with region A30-L34, i.e. within and adjacent to the MHR. The A$\beta$42 to A$\beta$40 difference map reveals pronounced isoform-specific differences in the DMPC/CHOL system (Figure 5I). The top five A$\beta$42-favored difference contacts are: R5-V18, D7-K16, L17-I32, F19-A30, and L34-V40. In contrast, the top five A$\beta$40-favored difference contacts are: G9-I32, E11-F20, G9-E22, E11-L34, and Y10-G33. Thus, in the DMPC/CHOL system, A$\beta$40 favors the tertiary contact network centered on region G9-E11, whereas A$\beta$42 favors contacts between the NTR and the CHC, in addition to contacts between the CHC and the MHR, and between the MHR and the CTR.

To determine how DMPC redistributes A$\beta$ tertiary contacts relative to lipid-free conditions, we analyzed the DMPC to no-lipids difference maps. In A$\beta$40, the top five DMPC-favored difference contacts are: F20-G29, E11-K16, F20-I31, E22-N27, and F20-A30 (Figure 5J). In contrast, the top five no-lipids-favored difference contacts are: E11-I32, G9-A30, G9-I31, Y10-I31, and V18-M35. Thus, the presence of DMPC shifts the tertiary organization of A$\beta$40 from contacts connecting regions G9-E11 and A30-I32 toward contacts connecting F20 with region G29-I31. In A$\beta$42, the top five DMPC-favored difference contacts are: E11-K16, L34-V40, V12-L17, G33-A42, and E11-V18 (Figure 5K). In contrast, the top five no-lipids-favored difference contacts are: L17-I32, F19-A30, H14-L34, H14-G33, and V18-I32. These difference maps thus indicate that the presence of DMPC lipids weakens contacts between regions H14-F19 and A30-L34 and rather enhances contacts E11-K16, E11-V18, V12-L17, and contacts between regions G33-L34 and the CTR.

To determine how the presence of DMPC lipids with CHOL redistributes A$\beta$ tertiary contacts relative to lipid-free conditions, we analyzed the DMPC/CHOL minus no-lipids difference contacts. In A$\beta$40, the top five DMPC/CHOL-favored difference contacts are: E11-V18, G9-F20, G9-G33, F20-G25, and F20-A30 (Figure 5L). In contrast, the top five no-lipids-favored difference contacts are: V18-M35, L17-V36, K16-G33, F20-I32, and H13-G33. Thus, in the presence of DMPC lipids with CHOL weakens contacts connecting region H13-F20 with region I32-V36 while favoring contacts between regions G9-E11 and V18-F20, between region G9-E11 and G33, alongside contacts F20-G25 and F20-A30. In A$\beta$42, the top five DMPC/CHOL-favored difference contacts are: D7-K16, R5-V18, E3-A21, E11-V18, and H6-G33 (Figure 5M). In contrast, the top five no-lipids-favored difference contacts are: F20-A30, F20-G29, F19-A30, E22-K28, and G9-G33. These differences indicate that the presence of DMPC/CHOL mixture shifts the tertiary organization of A$\beta$42 away from contacts between regions F19-E22 and K28-A30 and contact G9-G33 toward contacts connecting region E3-E11 with region K16-A21, alongside contact H6-G33. Taken together, these results reveal that lipids redistribute tertiary contacts rather than uniformly enhance or diminish them, whereby the resulting tertiary contact patterns depend on both lipid composition and A$\beta$ isoform.

#### Effect of lipids on the quaternary structure of A$\beta$ hexamers

To determine how A$\beta$ co-assembly with lipids affects quaternary structure of resulting A$\beta$40 and A$\beta$42 hexamers, we calculated intermolecular $C_\alpha$ contact maps for all three conditions: in the absence of lipids, in DMPC lipids, and in DMPC lipids mixed with CHOL.

In the absence of lipids, A$\beta$40 forms prominent intermolecular contacts between G25 and region G25-G33 (Figure 6A). Additional quaternary contacts are observed between G9 and region G33-G37, between regions K16-F20 and G33-V39, between regions G25-G29 and G37-G38, and between regions A2-H6 and G33-G37. The top five most probable quaternary contacts are: L17-L17, H13-G29, G9-G33, G25-N27, and G9-L34. The most probable contact L17-L17 facilitates proximity among pairs of CHCs. A broad distribution of quaternary contacts along the sequence indicates that the quaternary organization of A$\beta$40 is further supported by interactions between regions G25-G29 and G33-V39 as well as interactions between the NTR and the CHC. On the other hand, A$\beta$42 quaternary structure is characterized by prominent contacts between pairs of G29-A30 regions, centered around the most probable G29-G29 contact (Figure 6B). Additional quaternary contacts form between regions D7-G9 and G9-Y10 and between regions H6-H13 and G37-G38. The top five most probable contacts are: G29-G29, M35-M35, G29-A30, A30-G37, and K28-G29. The predominance of the above approximately in-register quaternary contacts indicates that A$\beta$42 favors more aligned in-parallel intermolecular contacts than A$\beta$40. This observation is further supported by the A$\beta$42 to A$\beta$40 difference contact map in Figure 6C. The top five A$\beta$40-favored difference contacts are L17-F19, H13-I31, G25-G29, G9-V36, and E11-G33. In contrast, the top five A$\beta$42-favored difference contacts are: G29-G29, M35-M35, G29-A30, K28-G29, and D7-G38, with the top four A$\beta$42-favored contacts support parallel in-register alignment of different peptides (Figure 6C). Thus, in the absence of lipids, quaternary structure of A$\beta$40 hexamers is characterized by contacts between pairs of CHC regions balance by contacts connecting G9, E11 and H13 with the MHR, whereas A$\beta$42 hexamers favor more parallel in-register arrangement.

Figure 6D shows that in the presence of DMPC lipids, the overall number of intermolecular contacts in A$\beta$40 hexamers is reduced, except for the contacts between pairs of the MHRs, which increase in the number. Thus, DMPC lipids substantially alter the quaternary structure of A$\beta$40 hexamers. The top five most probable intermolecular contacts are: K16-A30, A30-G33, H14-G33, K28-G37, and I31-G37. These contacts are located within regions H14-K16, K28-I31, and A30-G37, revealing that DMPC lipids favor an A$\beta$40 quaternary arrangement in which these A$\beta$40 regions are in close proximity. The presence of DMPC lipids reduces the number of intermolecular contacts in A$\beta$42 hexamers to a lesser degree, however, the overall contact probabilities are somewhat lower than in the absence of lipids. In the DMPC system, A$\beta$42 hexamers are stabilized by a cluster of intermolecular contacts between regions K16-F19 and M35-I41 as well as contacts between regions F20-V24 and G25-N27 (Figure 6E). The top five most probable contacts are: F20-A30, L17-I32, E22-K28, G33-G38, and I31-V40. Thus, the quaternary organization of A$\beta$42 hexamers involves rather heterogeneous and relatively low probability contacts between the CHC and the MHR, including residues adjacent to these regions, as well as interactions between the MHR and the CTR. The A$\beta$42 to A$\beta$40 difference contact map reveals distinct residue-specific contact preferences of the two isoforms (Figure 6F). The top five A$\beta$42-favored difference contacts are: F20-A30, L17-I32, E22-K28, G37-A42, and A21-I32, whereas the top five A$\beta$40-favored difference contacts are: K16-I32, H14-M35, K28-V39, I31-V39, and G29-V39. Thus, in the DMPC system, A$\beta$40 hexamers are stabilized by intermolecular contacts between regions H14-K16 and K28-I31 as well as the contacts between the MHR and the CTR, whereas A$\beta$42 hexamers are held together by intermolecular contacts between the CHC and region E22-A32 as well as contact G37-A42.

Figure 6G indicates that A$\beta$40 hexamers in the DMPC/CHOL system are held together by intermolecular contacts between regions G37-G38 and I31-G38 and between regions H14-G29 with G37-V39, in addition to a smaller localized cluster of contacts between regions A2-R5 and R5-S8. The top five most probable contacts are: F4-R5, G33-G38, F19-G37, G33-G37, and Q15-A21. On the other hand, A$\beta$42 hexamers are stabilized by parallel in-register intermolecular contacts along residues G33-G38 of pairs of peptides, alongside contacts connecting regions G37-V39 and V40-A42 (Figure 6H). Additional contributions to A$\beta$42 hexamer stability arise from a cluster of antiparallel strand-like intermolecular contacts connecting D1-H6 with G37-A42. A particularly strong same-residue intermolecular contact also forms between K16 and K16. The top five most probable intermolecular contacts in A$\beta$42 hexamers are: K16-K16, G33-G33, M35-M35, G37-G37, and G33-G37, indicating that the DMPC/CHOL system induces a parallel in-register alignment of neighboring peptides in A$\beta$42 hexamers that spans the CHC and the MHR. Consistent with these isoform-specific effects of lipids on the quaternary structure, the A$\beta$42 to A$\beta$40 difference contact map shows that contacts K16-K16, G33-G33, M35-M35, G37-G37, as well as H14-V18 are stronger in A$\beta$42 than in A$\beta$40. On the other hand, contacts F4-D7, G33-V40, F19-V39, G33-V39, and Q15-D23 are more probable in A$\beta$40 than in A$\beta$42 hexamers (Figure 6I). Thus, DMPC/CHOL promotes broadly distributed intermolecular contacts in A$\beta$40 hexamers but favors more ordered, aligned contacts involving in A$\beta$42 hexamers.

To investigate how DMPC lipids alter quaternary structure of A$\beta$ hexamers relative to lipid-free conditions, we analyzed the DMPC to no-lipids difference contact maps. In A$\beta$40 hexamers, the top five DMPC-favored difference contacts are: K16-A30, A30-G33, V12-F19, I31-G37, and H14-I32 (Figure 6J), whereas in A$\beta$42 hexamers, the top five no-lipids-favored difference contacts are: L17-L17, G9-G33, H13-G29, G25-N27, and G9-L34. Thus, DMPC lipids shift A$\beta$40 hexamer quaternary organization away from the same-residue L17 contact and the contact between G9 and H13 toward stronger individual intermolecular contacts between K16 and A30, A30 and G33, H14 and I32, and I31 and G37. In A$\beta$42 hexamers, the top five DMPC-favored difference contacts are: F20-A30, L17-I32, E22-K28, I31-V40, and G33-G38, whereas the top five no-lipids-favored difference contacts are: G29-G29, M35-M35, G29-A30, K28-G29, and D7-G38 (Figure 6K). Thus, in A$\beta$42 hexamers, DMPC lipids inhibit intermolecular contacts G29-G29 and M35-35 and instead promote cross-region contacts involving the CHC, E22-K28, and residues within or adjacent to the MHR and the CTR.

Figure 6L-M with the lipids to no-lipids difference contact maps reveals distinct changes in the quaternary organization in hexamers of the two isoforms in the DMPC/CHOL system. In A$\beta$40 hexamers, the top five largest lipids-favored difference contacts are: F4-R5, F19-G37, Q15-A21, A2-D7, and R5-G33, whereas the top five no lipids-favored difference contacts are: G9-G33, H13-G29, L17-L17, G25-N27, and S26-G37 (Figure 6L). The system of DMPC lipids mixed with CHOL shifts quaternary contacts in A$\beta$40 hexamers away from contacts between regions G9-H13 and G29-G33 and contact L17-L17 toward contacts A2-R5, Q15-A21, F19-G37, and R5-G33, which involve the contacts within the NTR and between the NTR and the MHR. A$\beta$40 also forms a parallel strand-like series of intermolecular contacts connecting Y10-Q15 with K16-A21 in the presence of DMPC/CHOL lipids. In A$\beta$42 hexamers, the top five most probable lipids-favored difference contacts are: K16-K16, G33-G33, G37-G37, G33-G37, and H14-V18, whereas the top five no lipids-favored difference contacts are G29-G29, G29-A30, K28-G29, A30-G37, and D7-G38 (Figure 6M). Thus, DMPC lipids with CHOL weaken quaternary contacts in A$\beta$42 hexamers that are centered around contact K28-A30 and instead promote parallel in-register aligned same-residue contacts.

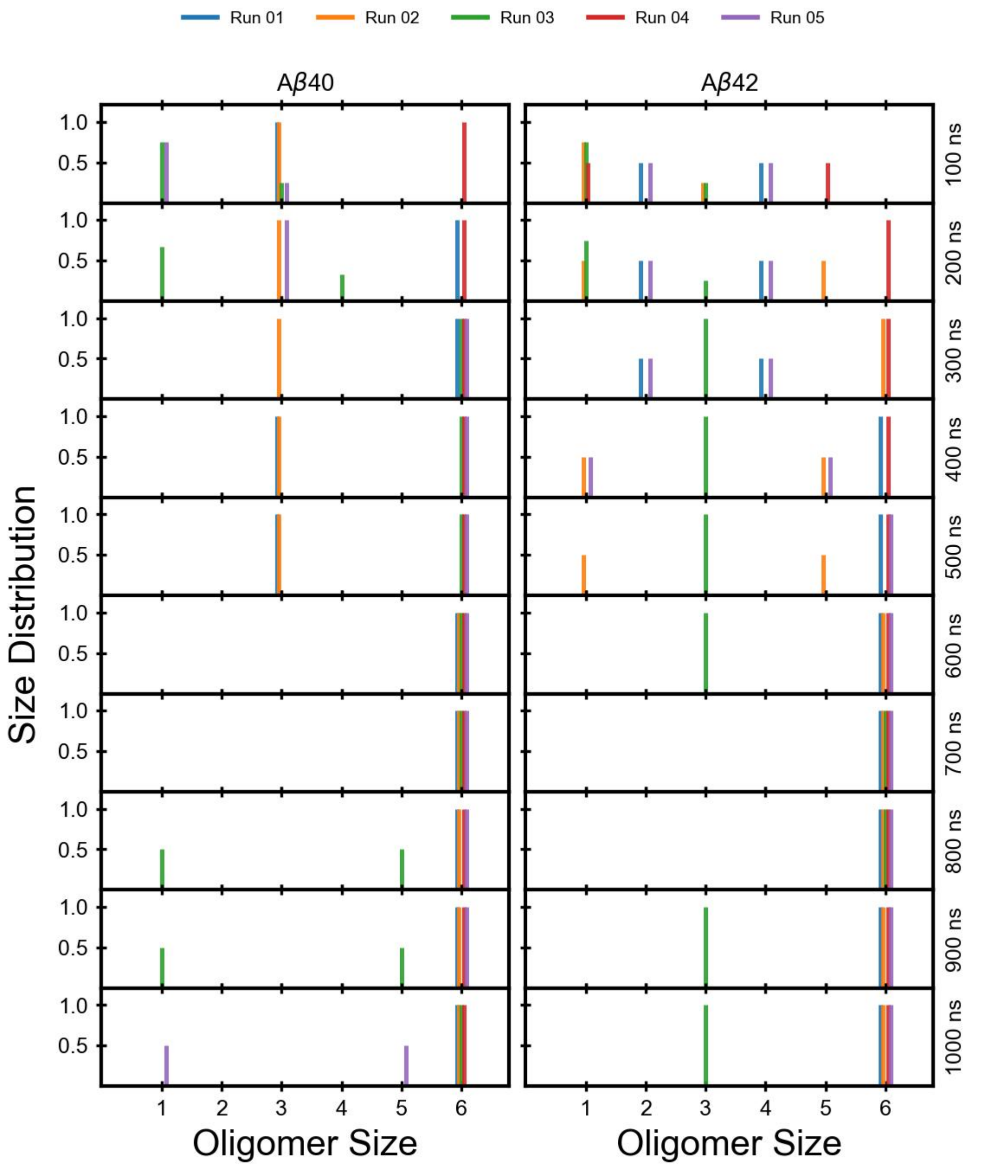


**Fig. S1** Time evolution of A$\beta$40 and A$\beta$42 oligomer size distribution for all five trajectories. The size distributions are calculated for each trajectory individually every 100 ns, starting from 100 ns.

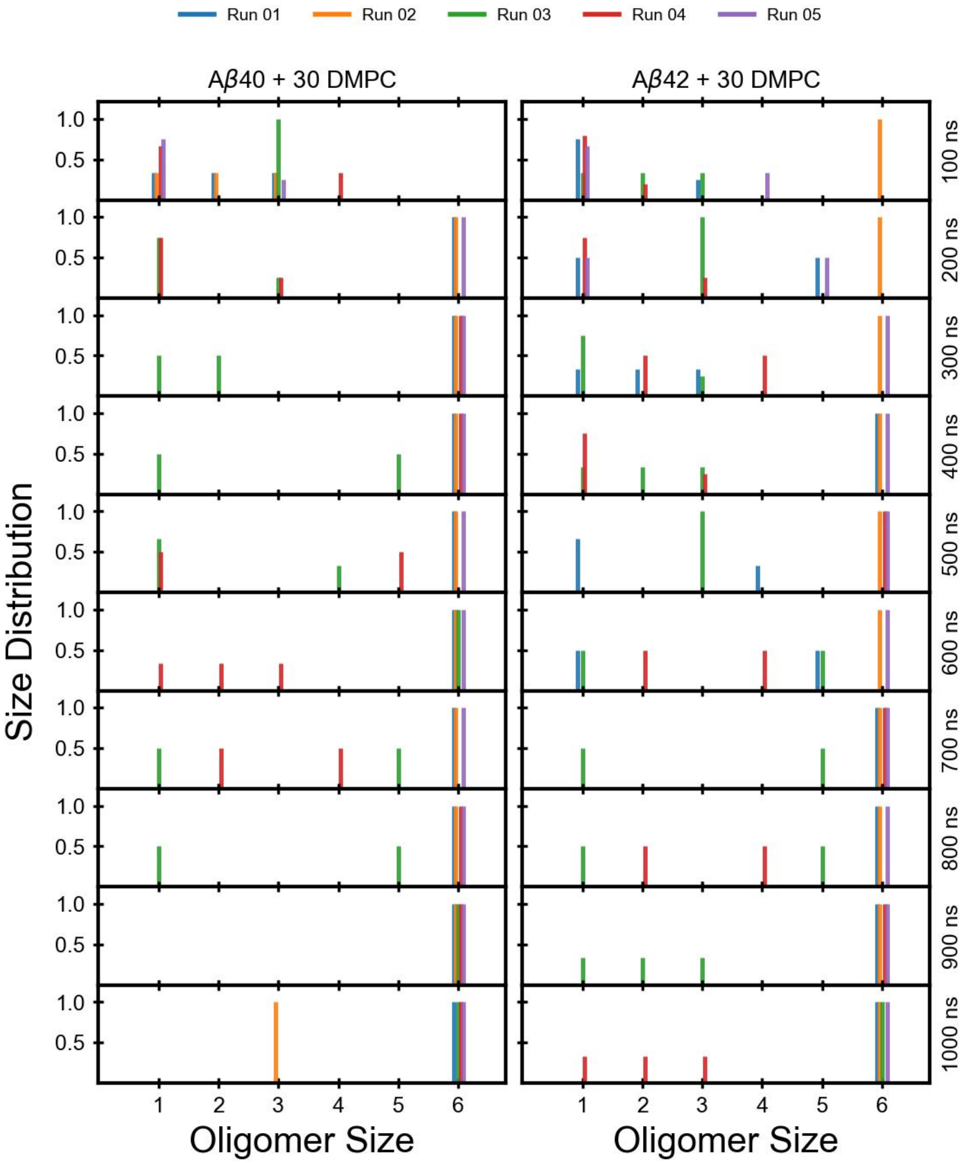


**Fig. S2** Time evolution of A$\beta$40 and A$\beta$42 oligomer size distributions in the presence of 30 DMPC lipids for all five trajectories. The size distributions are calculated for each trajectory individually every 100 ns, starting from 100 ns.

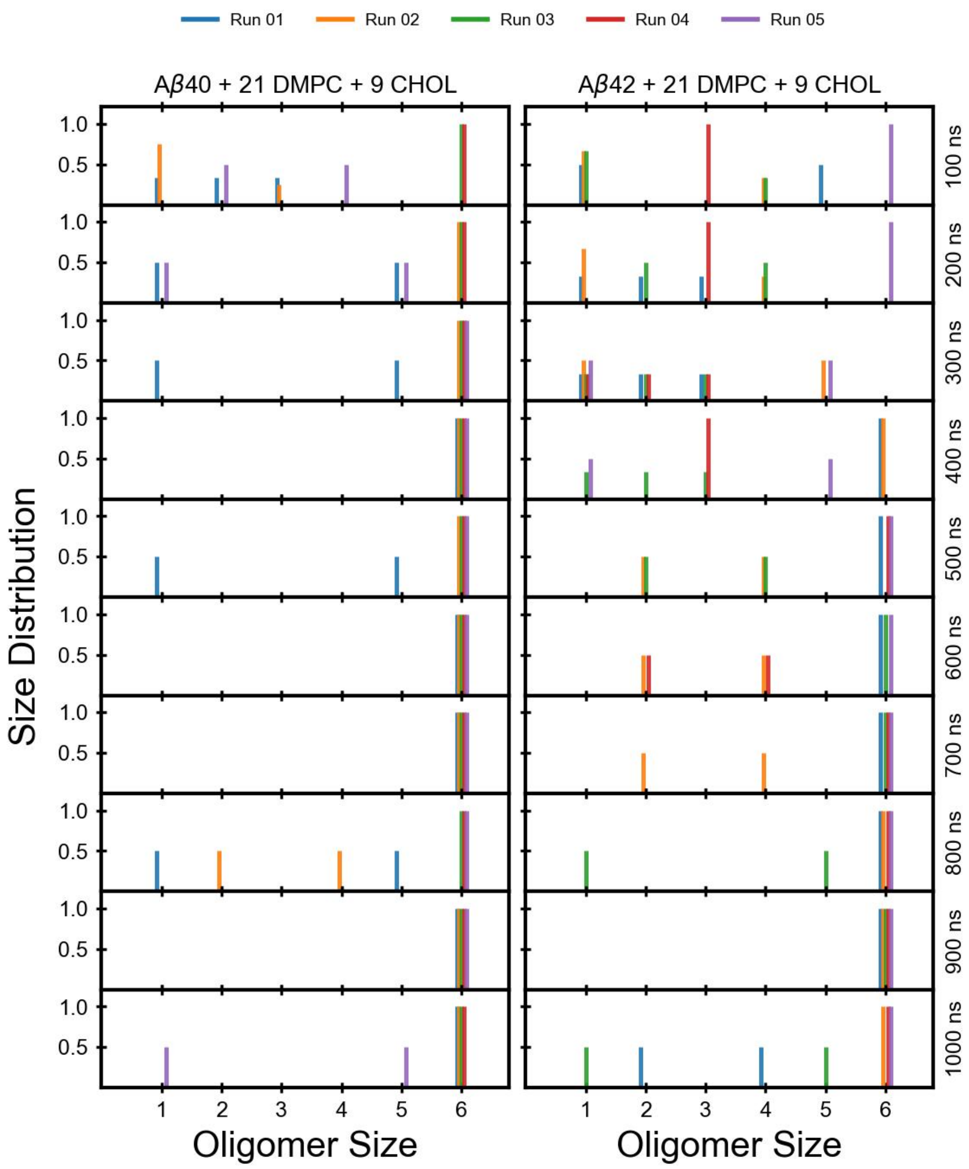


**Fig. S3** Time evolution of A$\beta$40 and A$\beta$42 oligomer size distributions in the presence of 21 DMPC and 9 CHOL lipids for all five trajectories. The size distributions are calculated for each trajectory individually every 100 ns, starting from 100 ns.

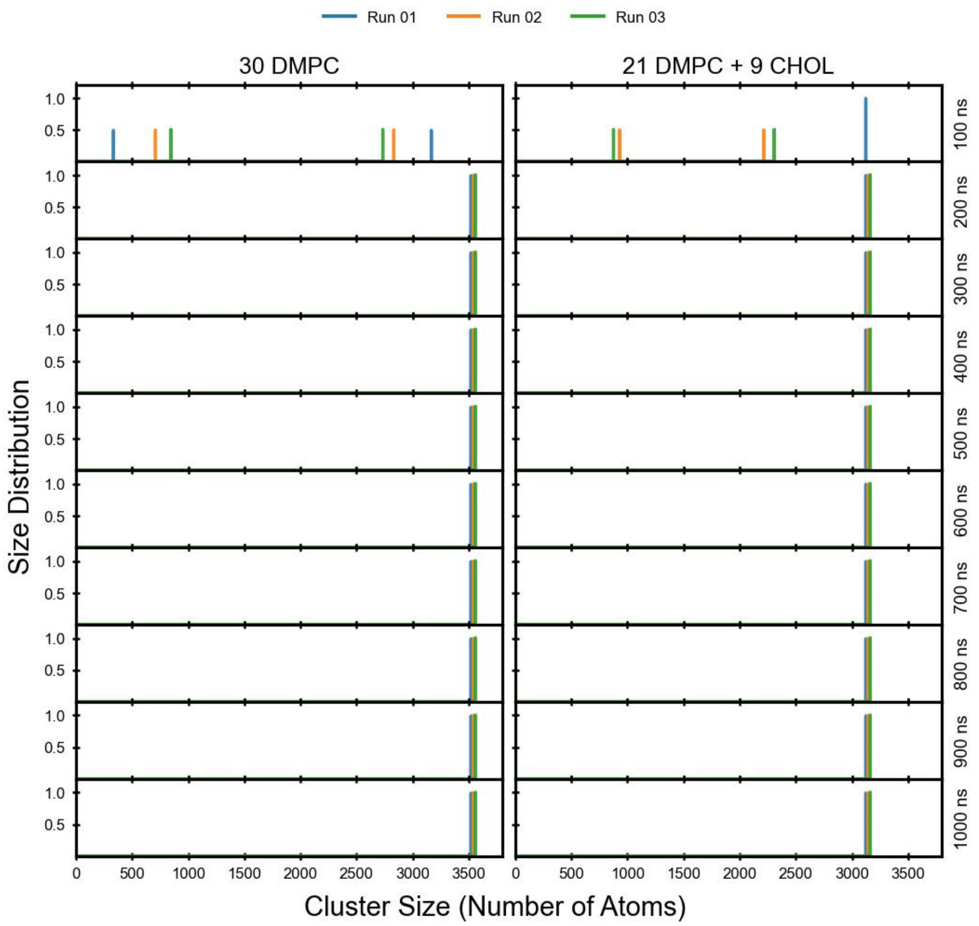


**Fig. S4** Time evolution of cluster size distributions for the lipids in two systems (i) 30 DMPC, (ii) 21 DMPC and 9 CHOL molecules

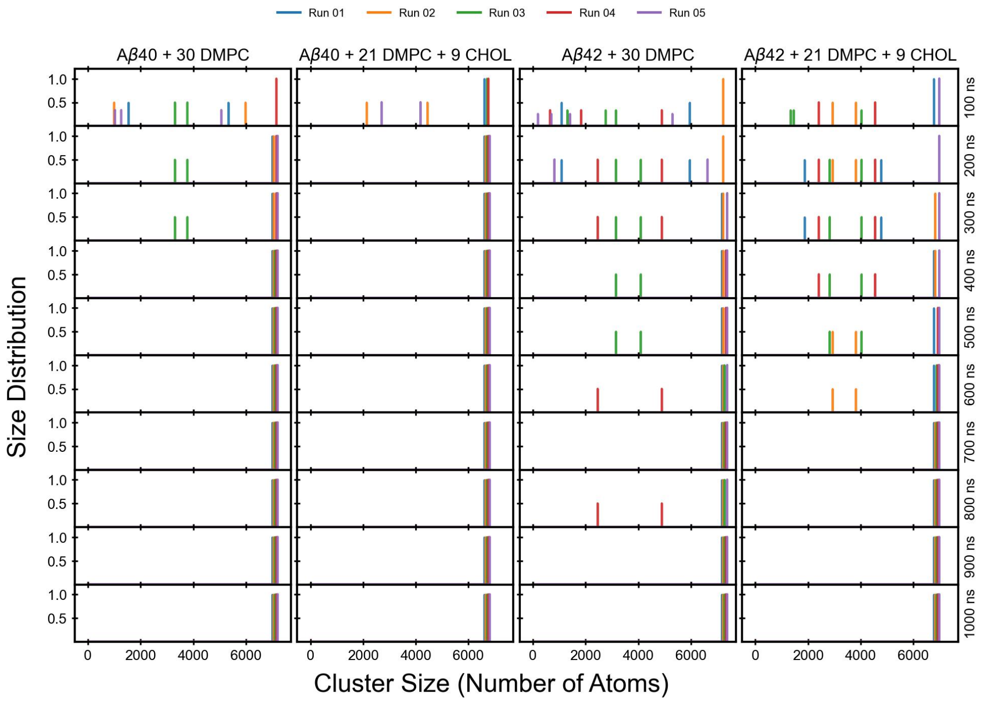


**Fig. S5** Time evolution of the total cluster size distributions of Aβ40 and Aβ42 in two systems (i) with 30 DMPC, and (ii) with 21 DMPC and 9 CHOL molecules. The distributions are calculated every 100 ns for all the trajectories starting from 100 ns.

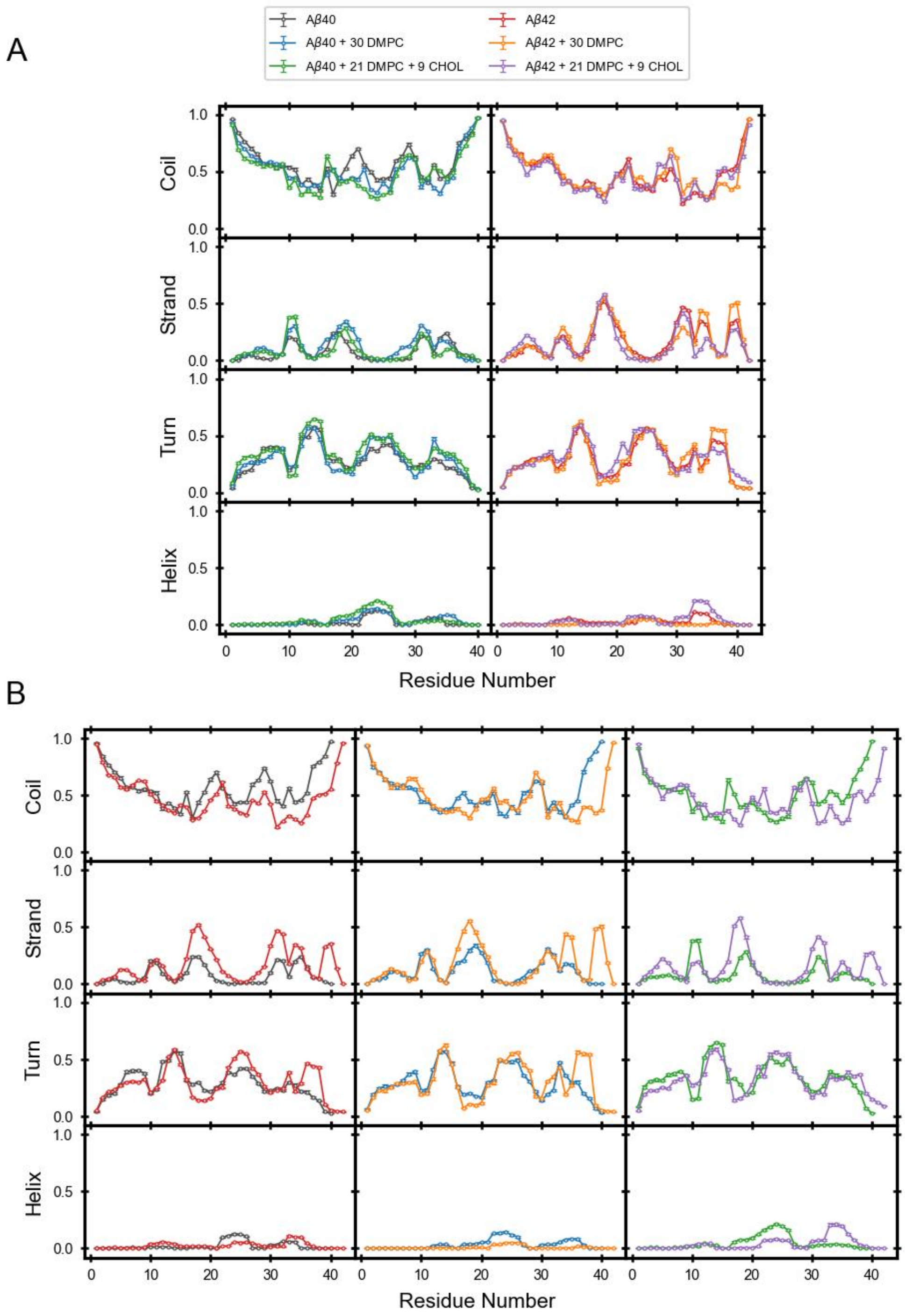


**Fig. S6** Per-residue secondary structure propensities of Aβ40 and Aβ42 (i) without lipids, (ii) with 30 DMPC, (iii) with 21 DMPC and 9 CHOL

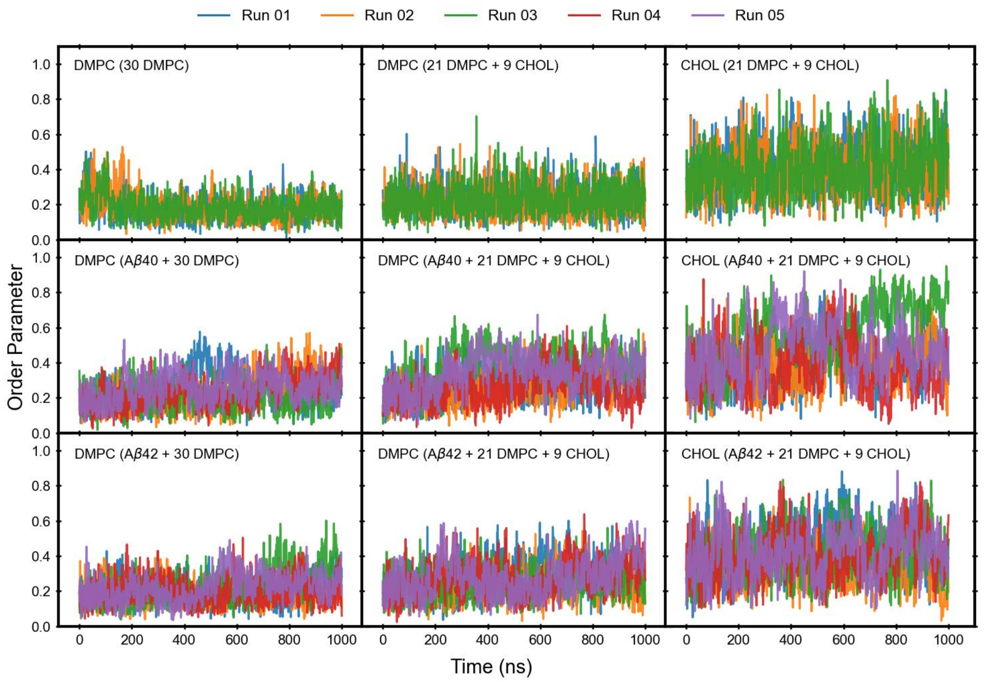


**Fig. S7** Time evolution of the nematic order parameter, $S_2$, of DMPC and CHOL molecules in the lipids-only systems (top row), systems containing A$\beta$40 (middle row), and systems containing A$\beta$42 (bottom row). The left column shows DMPC in the DMPC system, whereas the middle and right columns show DMPC and CHOL, respectively, in the DMPC/CHOL system. Three replica trajectories are shown for the lipids-only systems and five replica trajectories for the A$\beta$40 and A$\beta$42 systems, with individual trajectories represented by distinct colors.

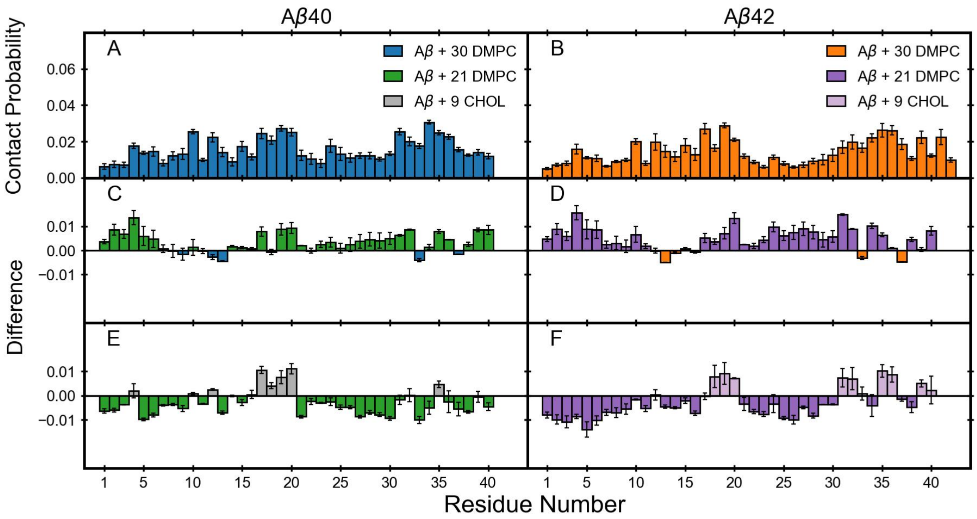


**Fig. S8** Per-residue contact probabilities of DMPC with (A) A$\beta$40 and (B) A$\beta$42 in the DMPC system. (C,D) Differences in A$\beta$-DMPC contact probabilities between the DMPC/CHOL and DMPC systems, calculated as the contact probabilities with 21 DMPC in the DMPC/CHOL system minus those with 30 DMPC in the DMPC system, for (C) A$\beta$40 and (D) A$\beta$42. (E,F) Differences between A$\beta$-CHOL contact probabilities in the DMPC/CHOL system and A$\beta$-DMPC contact probabilities in the DMPC system, calculated as the contact probabilities with 9 CHOL minus those with 30 DMPC, for (E) A$\beta$40 and (F) A$\beta$42. The final values are averaged over 600-1000 ns and across all trajectories, with error bars representing the SEM.

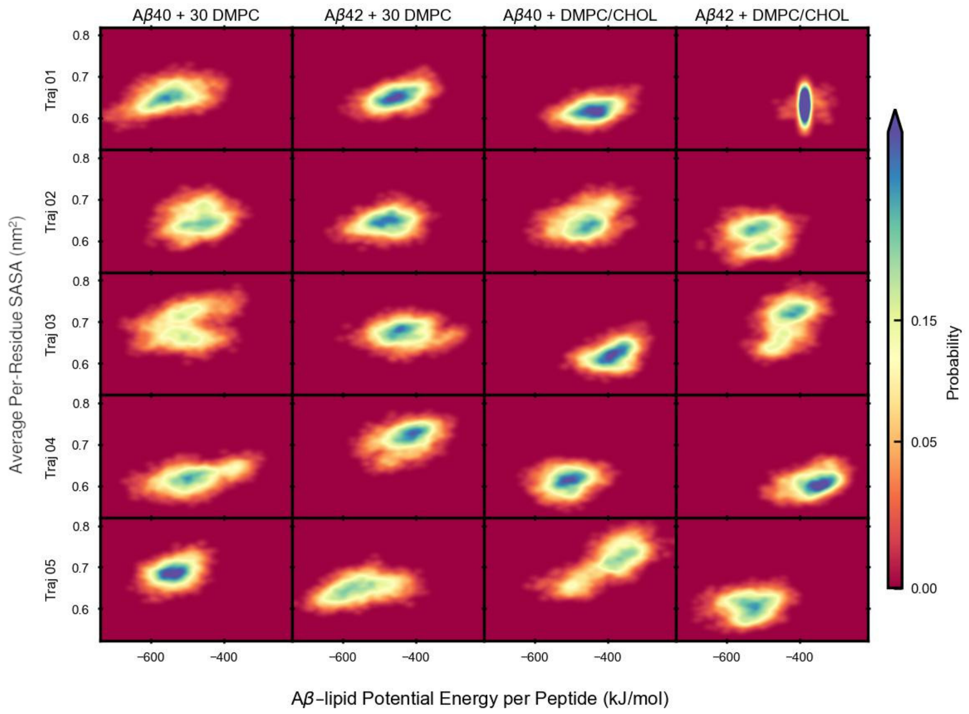


**Fig. S9** Two-dimensional probability distributions of the A$\beta$-lipid potential energy per peptide and average solvent-accessible surface area (SASA) per residue for individual trajectories. The four columns correspond to A$\beta$40 with DMPC, A$\beta$42 with DMPC, A$\beta$40 with DMPC/CHOL, and A$\beta$42 with DMPC/CHOL, respectively. Each row represents one of the five replica trajectories for the corresponding system. All analyses were performed over 900-1000 ns.
